# Integrated Multiomics Reveals Intergenerational Brain Sensory System Remodeling Induced by Parental Intermittent Fasting

**DOI:** 10.64898/2026.10.15.759492

**Authors:** Xiangyuan Peng, Vernise JT Lim, Yibo Fan, Nishat I. Tabassum, Xiangru Cheng, Marconi Fung, Grant R. Drummond, Christopher G. Sobey, Bushra Amin, Rohan Lowe, Yong U. Liu, Mitchell K. P. Lai, Guobing Chen, Mark P. Mattson, Dong-Gyu Jo, Terrance G. Johns, Oscar Junhong Luo, Thiruma V. Arumugam

**Author notes:** Corresponding authors: Thiruma V. Arumugam or Xiangyuan Peng, La Trobe Institute for Molecular Science and Department of Microbiology, Anatomy, Physiology and Pharmacology, School of Agriculture, Biomedicine and Environment, La Trobe University, Bundoora, VIC, Australia.; or. **Author contributions:** Study conception and design: T.V.A., X.P., T.G.J., M.P.M., D.G.J. and O.J.L.; experiment or data collection: X.P., Y.F., V.J.T.L., N.I.T., X.C., M.C., B.A., R.L., and T.V.A.: data analysis: X.P., V.J.T.L., Y.F., and T.V.A.; data analytics and interpretation: X.P., V.J.T.L., Y.F., O.J.L., and T.V.A.; writing-manuscript preparation and intellectual input: X.P., G.R.D., C.G.S., Y.U.L., M.K.P.L., G.C., M.P.M., D.G.J., T.G.J., O.J.L., and T.V.A.; supervision and administration: T.V.A. **Competing Interest Statement:** T.G.J. is affiliated with Epigenes Pty Ltd. The remaining authors declare no competing interests.

## Abstract

Intermittent fasting (IF) is increasingly recognized as a potent modulator of metabolic health, but its effects on brain function across generations remain poorly understood. Here, we show that parental IF, administered to both mothers and fathers before mating, is associated with coordinated molecular remodeling in the cerebral cortex of fathers and their offspring, with shared molecular signatures identified between these generations and a pronounced enrichment of sensory-associated pathways. Using whole-genome bisulfite sequencing, transcriptomic profiling, and proteomic analysis, we found that IF elicited widespread cortical epigenetic remodeling in fathers, accompanied by transcriptional and protein-level changes enriched for olfactory, taste, synaptic, and neurometabolic pathways. Strikingly, related molecular signatures were detected in F1 offspring that had never been directly exposed to fasting, consistent with intergenerational transmission of IF-associated cortical remodeling. Offspring responses were sex dependent, with female and male progeny exhibiting distinct patterns of DNA methylation, gene expression, and protein abundance. Integrated multiomic analyses identified conserved candidate networks involved in sensory perception, glutamatergic signaling, synaptic organization, intracellular trafficking, cytoskeletal regulation, and stress-response pathways. Together, these findings identify sensory-associated cortical remodeling as a prominent molecular feature associated with parental IF and reveal persistent, sex-dependent molecular alterations in the offspring brain.

**Significance Statement:** This study shows that intermittent fasting induces intergenerational remodeling of the cerebral cortex across epigenetic, transcriptomic, and proteomic layers. By integrating multiomic profiling in fathers and unexposed offspring, we found that intermittent fasting prominently alters sensory-associated pathways, including olfactory, taste, glutamatergic, and synaptic signaling networks. These effects were conserved across generations and displayed marked sex-specific patterns in offspring cortex. Our findings identify sensory neurobiology as a previously unrecognized target of intergenerational dietary programming and suggest that parental feeding patterns can shape descendant brain molecular architecture.

## Introduction

Over the past several decades, intermittent fasting (IF) has emerged as a powerful non-pharmacological intervention capable of improving healthspan, metabolic fitness, and cellular resilience. Unlike conventional dietary restriction paradigms, IF involves periodic fasting or restricting food intake to defined temporal windows without necessarily reducing total caloric intake. Accumulating evidence demonstrates that these feeding paradigms mitigate age-associated cardiovascular, neurodegenerative, inflammatory, and metabolic disorders while enhancing cognitive performance, stress resistance, and longevity across multiple species [1–4]. IF is increasingly recognized as a potent regulator of molecular plasticity within peripheral tissues and the central nervous system.

Previous work showed that IF induces both shared and tissue-specific adaptive responses across heart, liver, skeletal muscle, and brain [5, 6], reprograms the brain gene expression and proteome to protect against neuroinflammation, synaptic degeneration and cognitive impairment in stroke and vascular dementia (VaD) [7–11], and remodels the DNA methylation landscape while attenuating hallmark neuropathology in experimental VaD [8]. Moreover, IF alters histone trimethylation dynamics and transcriptional programs associated with metabolic adaptation [12], and that global DNA methylation within CG-rich nucleotide islands is extensively remodeled under IF conditions [8]. Furthermore, IF induces widespread chromatin accessibility remodeling linked to coordinated transcriptional reprogramming and IF induces widespread, tissue-specific reprogramming of transposable elements which in turn influence the expression of genes in a non-random manner to reprogram cellular physiology [13, 14]. Collectively, these findings establish meal timing and frequency affect cellular resilience operating via coordinated metabolic, epigenetic, transcriptomic, and proteomic reprogramming.

Epigenetic regulation has emerged as a central mechanism through which environmental experiences become biologically embedded to shape long-term phenotypes. DNA methylation, histone modifications, chromatin remodeling, and non-coding RNA activity collectively form a dynamic regulatory architecture capable of translating environmental cues into stable transcriptional states. Importantly, growing evidence suggests that environmentally induced epigenetic changes may extend beyond directly exposed individuals and influence subsequent generations [15–18]. Such inter- and transgenerational inheritance has profound implications for understanding how environments shape offspring physiology, disease susceptibility, and neural function. To date, studies investigating intergenerational effects of dietary interventions have focused predominantly on maternal caloric restriction or nutrient imbalance. Within the context of IF, maternal alternate-day fasting has been reported to alter hepatic DNA methylation, DNA methyltransferase expression, and mTORC1 signaling in offspring [19, 20]. Other studies have demonstrated that maternal IF prior to pregnancy can disrupt offspring intestinal homeostasis and predispose offspring to glucose and lipid metabolic abnormalities [21], while prolonged maternal fasting increased susceptibility to renal injury under salt challenge conditions in male offspring [22]. Although these studies provide important insights into developmental programming by maternal dietary interventions, several major questions remain unresolved. First, the impact of combined maternal and paternal IF, or paternal IF independently, on offspring biology remains largely unknown. Second, previous studies have focused almost exclusively on metabolic phenotypes, leaving higher-order neural and sensory functions unexplored. Third, no prior study has comprehensively integrated epigenomic, transcriptomic, and proteomic approaches to identify the molecular architecture underlying intergenerational transmission induced by dietary timing. We recently provided the first evidence that parental IF alters metabolic phenotypes in unexposed offspring, including body weight regulation, glucose tolerance, and metabolic gene expression [23]. These findings established proof-of-principle that parental feeding schedules before conception can transmit metabolic memory across generations. However, whether these intergenerational effects extend beyond metabolism into sensory neurobiology, and what molecular mechanisms govern such transmission, remained unknown.

Sensory processing is dynamically coupled to nutritional and metabolic state. Previous behavioral studies have shown that food restriction can alter olfactory behavior in rodents. In particular, food restriction influenced the expression of conditioned odor discrimination in mice, suggesting that nutritional state can modulate olfactory responses and odor-guided behavior [24]. Consistent with this concept, fasting has been shown to alter olfactory neural processing, including increased spontaneous and odor-responsive activity in olfactory-bulb mitral/tufted cells and changes in odor representation, indicating that metabolic state can directly influence sensory circuit function [25]. The interaction between nutritional state and olfactory processing is further supported by studies demonstrating that metabolic signals regulate olfactory pathways involved in food-seeking and feeding behavior [26]. More recent work also suggests that caloric restriction and intermittent fasting can influence peripheral taste-related features, indicating that dietary restriction may affect multiple components of chemosensory biology [27]. However, the molecular mechanisms through which sustained dietary restriction or time-restricted feeding remodels sensory systems remain poorly understood, particularly at the level of integrated epigenetic, transcriptional, and proteomic regulation. Thus, the sensory-associated molecular signatures identified in the present study may represent a molecular substrate through which altered nutritional state influences sensory biology, although direct functional consequences for sensory performance remain to be established. Therefore, in the present study, we investigated whether parental IF induces intergenerational sensory remodeling and sought to define the integrated molecular signatures underlying this phenomenon. To address this, we performed whole-genome bisulfite sequencing (WGBS), RNA sequencing, and proteomic profiling across sensory-associated brain regions in both parental and offspring generations. Using this integrated multiomics framework, we demonstrate for the first time that parental IF is associated with intergenerational remodeling of molecular pathways related to tactile, olfactory, and proprioceptive function in unexposed offspring. Moreover, we identify convergent epigenetic, transcriptomic, and proteomic remodeling involving synaptic organization, neuroimmune signaling, neurometabolic pathways, and sensory-associated gene networks that distinguish offspring of IF parents from controls. Collectively, these findings reveal that parental feeding schedules shape sensory neurobiology across generations and provide a comprehensive multiomic framework for understanding how ancestral dietary timing reprograms neural information processing in descendants.

## Results

### Intermittent fasting reprograms the cortical methylome and sensory-associated pathways in paternal mice

Male and female C57BL/6 mice were subjected to either ad libitum (AL) feeding or a 16-hour daily IF regimen for four months prior to mating (**Fig. 1A**). Under the IF paradigm, animals underwent fasting from 4:00 pm to 8:00 am with an 8-hour feeding window between 8:00 am and 4:00 pm, while AL mice had unrestricted access to food throughout the study. After completion of the dietary intervention, animals from the same dietary groups were mated (AL♂ × AL♀ and IF♂ × IF♀) to generate offspring. Following weaning, all offspring were maintained under standard AL feeding conditions for four months prior to tissue collection, thereby excluding direct fasting exposure in the offspring generation. Throughout the experimental period, body weight and blood metabolic parameters were monitored. At the endpoint, cerebral cortex tissue was collected for integrated multiomic analyses, including whole-genome bisulfite sequencing (WGBS), RNA sequencing, and proteomic profiling, to investigate IF-associated molecular remodeling in fathers and both female and male offspring. The overall experimental workflow included genome-wide DNA methylation analysis, transcriptomic profiling, proteomic analysis, and downstream pathway enrichment analyses. Male mice subjected to a 16-hour daily IF regimen for four months exhibited significant physiological adaptations compared with AL controls (**SI Appendix, Fig. S1**). Although both groups progressively gained body weight over time, IF mice displayed significantly attenuated weight gain beginning approximately at week 5 and continuing throughout the intervention period (**SI Appendix, Fig. S1 A**). Consistent with fasting-induced metabolic adaptation, blood glucose levels were significantly reduced in IF mice at 0, 8, and 16 weeks compared with AL controls (**SI Appendix, Fig. S1 B**). Conversely, circulating ketone levels were significantly elevated in IF mice, particularly at later time points (**SI Appendix, Fig. S1 C**), confirming effective induction of fasting-associated metabolic switching prior to tissue collection.

**Figure 1.**
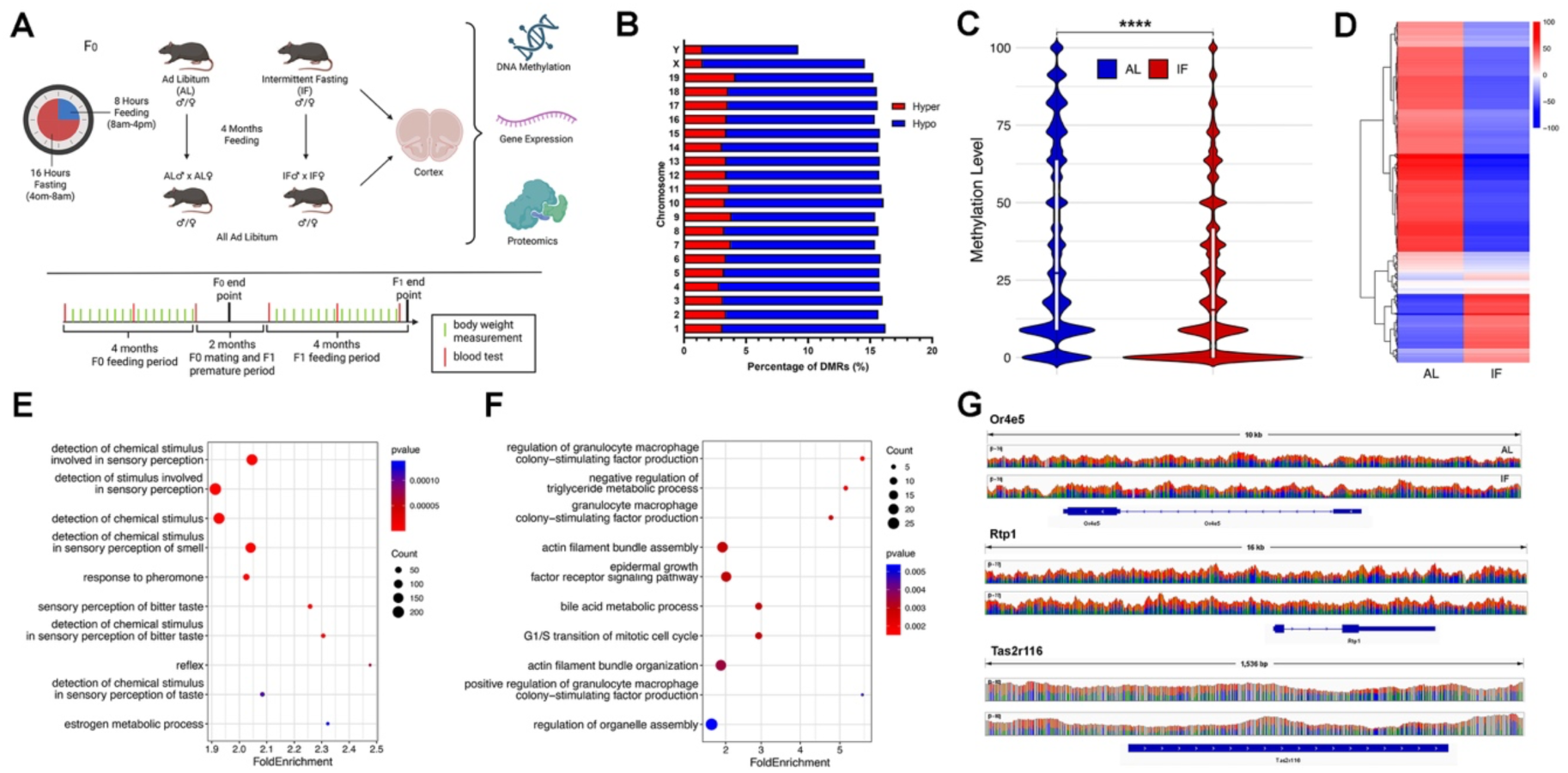
Intermittent fasting induces extensive sensory-associated cortical DNA methylation remodeling in paternal mice. (A) Experimental schematic illustrating the intermittent fasting (IF) paradigm and multiomic workflow. Male and female C57BL/6 mice were subjected to either ad libitum (AL) feeding or a 16-hour daily IF regimen for four months prior to mating. IF animals underwent fasting from 4:00 pm to 8:00 am with an 8-hour feeding window between 8:00 am and 4:00 pm. Following dietary intervention, animals were mated within dietary groups (AL♂ × AL♀ and IF♂ × IF♀). All offspring were subsequently maintained under standard AL feeding conditions for four months prior to tissue collection. Cerebral cortex tissue was collected for integrated multiomic analyses including whole-genome bisulfite sequencing (WGBS), RNA sequencing, and proteomic profiling. Body weight and blood metabolic parameters were monitored throughout the experimental period. (B) Chromosomal distribution of differentially methylated regions (DMRs) in paternal IF cortex samples relative to AL controls. Hypomethylated DMRs predominated across most chromosomes. (C) Violin plot showing global methylation level distributions between AL and IF paternal cortex samples. ****P < 0.0001. (D) Hierarchical clustering heatmap of differentially methylated genes demonstrating clear segregation between AL and IF cortical methylation profiles. (E) Gene ontology (GO) enrichment analysis of hypomethylated genes in IF cortex samples, showing enrichment of sensory-associated pathways including olfactory, taste, and pheromone signaling. (F) GO enrichment analysis of hypermethylated genes in IF cortex samples, showing enrichment of pathways associated with metabolic regulation, immune signaling, cytoskeletal organization, and organelle assembly. (G) Representative methylation tracks showing altered methylation landscapes across sensory-associated loci including *Or4e5, Rtp1,* and *Tas2r116* in IF cortex samples relative to AL controls.

Principal component analysis (PCA) of methylation profiles demonstrated clear separation between AL and IF cortical samples (**SI Appendix, Fig. S2**), indicating that IF induced distinct global methylation signatures within the paternal cerebral cortex. Samples from IF mice clustered independently from AL controls across principal component dimensions, further supporting substantial epigenomic divergence induced by prolonged IF exposure. Genome-wide analysis identified widespread differentially methylated regions (DMRs) between IF cortex and AL cortex distributed across all autosomes and sex chromosomes (**Fig. 1B**). Across most chromosomes, compared with the AL group, the IF group exhibited a predominance of hypomethylated DMRs over hypermethylated regions, indicating that IF induced a broad shift toward cortical hypomethylation. Global methylation distribution analysis further demonstrated significant alterations in methylation levels between AL and IF groups (**Fig. 1C**). Hierarchical clustering analysis revealed clear segregation of methylation profiles according to dietary condition (**Fig. 1D**), indicating robust IF-induced epigenomic remodeling within the cerebral cortex. Heatmap analysis of differentially methylated genes with a >25% difference in methylation levels within promoter regions revealed extensive remodeling of sensory-associated gene families in response to IF. A large subset of olfactory receptor and taste receptor genes exhibited reduced methylation levels in IF cortex samples compared with AL cortex samples (**SI Appendix, Fig. S3 A**). These included multiple members of the *Olfr* and *Tas2r* gene families, together with sensory signaling-related genes such as *Gnat3* and *Trpa1*. Conversely, heatmap analysis of hypermethylated differentially methylated genes revealed increased methylation across a distinct subset of genes in IF cortex samples compared with AL cortex samples (**SI Appendix, Fig. S3 B**). These hypermethylated genes included regulators associated with neuronal signaling, cellular organization, metabolic adaptation, and stress-response pathways, including Epha10, Syn2, Hif1a, and Sirt1, indicating that IF induces selective epigenetic repression across multiple functional cortical networks.

Gene ontology (GO) enrichment analysis of differentially methylated genes demonstrated prominent enrichment of sensory-associated biological processes in IF cortex samples. Hypomethylated genes were significantly enriched in pathways including detection of chemical stimulus involved in sensory perception, sensory perception of smell, pheromone response, bitter taste perception, taste receptor signaling, and reflex-associated pathways (**Fig. 1E**). These findings indicate that IF strongly remodels methylation patterns in genes involved in chemosensory processing and environmental signal detection. In contrast, hypermethylated genes were enriched in pathways associated with granulocyte macrophage colony-stimulating factor regulation, triglyceride metabolic processes, epidermal growth factor receptor signaling, bile acid metabolism, G1/S transition of the mitotic cell cycle, actin filament organization, and organelle assembly (**Fig. 1F**), suggesting concurrent modulation of metabolic, immune, and cytoskeletal regulatory pathways under IF conditions. Representative methylation tracks further demonstrated altered methylation landscapes across sensory-associated loci including *Or4e5*, *Rtp1*, and *Tas2r116* in IF cortex samples relative to AL controls (**Fig. 1G**), confirming locus-specific methylation remodeling in genes linked to olfactory and taste signaling.

Further GO enrichment analysis of differentially methylated genes demonstrated substantial functional reorganization associated with IF-induced methylation changes. Among hypomethylated-associated pathways, enriched cellular component categories included GABAergic synapse, actin-based cell projections, filopodia, and intercellular canaliculi (**SI Appendix, Fig. S4 A**). Molecular function enrichment analysis identified odorant binding, pheromone receptor activity, taste receptor activity, prolactin receptor binding, and trace amine receptor activity as major affected pathways (**SI Appendix, Fig. S4 B**). In contrast, hypermethylated-associated pathways were enriched for peroxisome, microbody, mitotic spindle, ribosomal complexes, and synaptic vesicle membrane components (**SI Appendix, Fig. S4 C**). Molecular function categories additionally included phosphatase regulator activity, protein heterodimerization activity, ubiquitin ligase binding, and metal ion sensor activity (**SI Appendix, Fig. S4 D**). Kyoto Encyclopedia of Genes and Genomes (KEGG) pathway analysis further revealed significant enrichment of neuronal and metabolic signaling pathways among differentially methylated genes. Hypomethylated-associated pathways included glutamatergic synapse, long-term potentiation, longevity regulating pathway, glucagon signaling pathway, taste transduction, and spinocerebellar ataxia pathways (**SI Appendix, Fig. S5 A**). In contrast, hypermethylated-associated pathways included mismatch repair, nucleotide excision repair, pyruvate metabolism, fatty acid degradation, ubiquitin-mediated proteolysis, vascular smooth muscle contraction, and peroxisome pathways (**SI Appendix, Fig. S5 B**).

Collectively, these findings demonstrate that prolonged IF induces extensive cortical DNA methylation remodeling in paternal mice, prominently affecting sensory-associated signaling pathways together with neuronal, metabolic, and cellular regulatory networks.

### Integrated multiomic analysis reveals coordinated epigenetic, transcriptomic, and proteomic remodeling in the cortex of intermittent fasting fathers

To determine whether IF induces coordinated molecular alterations beyond DNA methylation remodeling, integrative analyses combining transcriptomic and proteomic datasets with WGBS signatures were performed using cerebral cortex tissue collected from F_0_ fathers. Heatmap analyses demonstrated clear segregation between AL and IF cortical samples across all molecular platforms, including DNA methylation (**Fig. 2A**), transcriptomic profiles (**Fig. 2B**), and proteomic signatures (**Fig. 2C**), indicating robust IF-induced remodeling at epigenetic, transcriptional, and protein levels. Pairwise correlation analysis revealed strong concordance among the sample-level component scores derived from the RNA-seq, proteomic, and DNA methylation datasets (**Fig. 2D**). The RNA-seq and proteomic components showed the strongest correlation (r = 0.99), followed by the correlations between RNA-seq and DNA methylation components (r = 0.82) and between proteomic and DNA methylation components (r = 0.86). These results indicate substantial concordance among the multiomic datasets in the paternal cerebral cortex under IF conditions.

**Figure 2.**
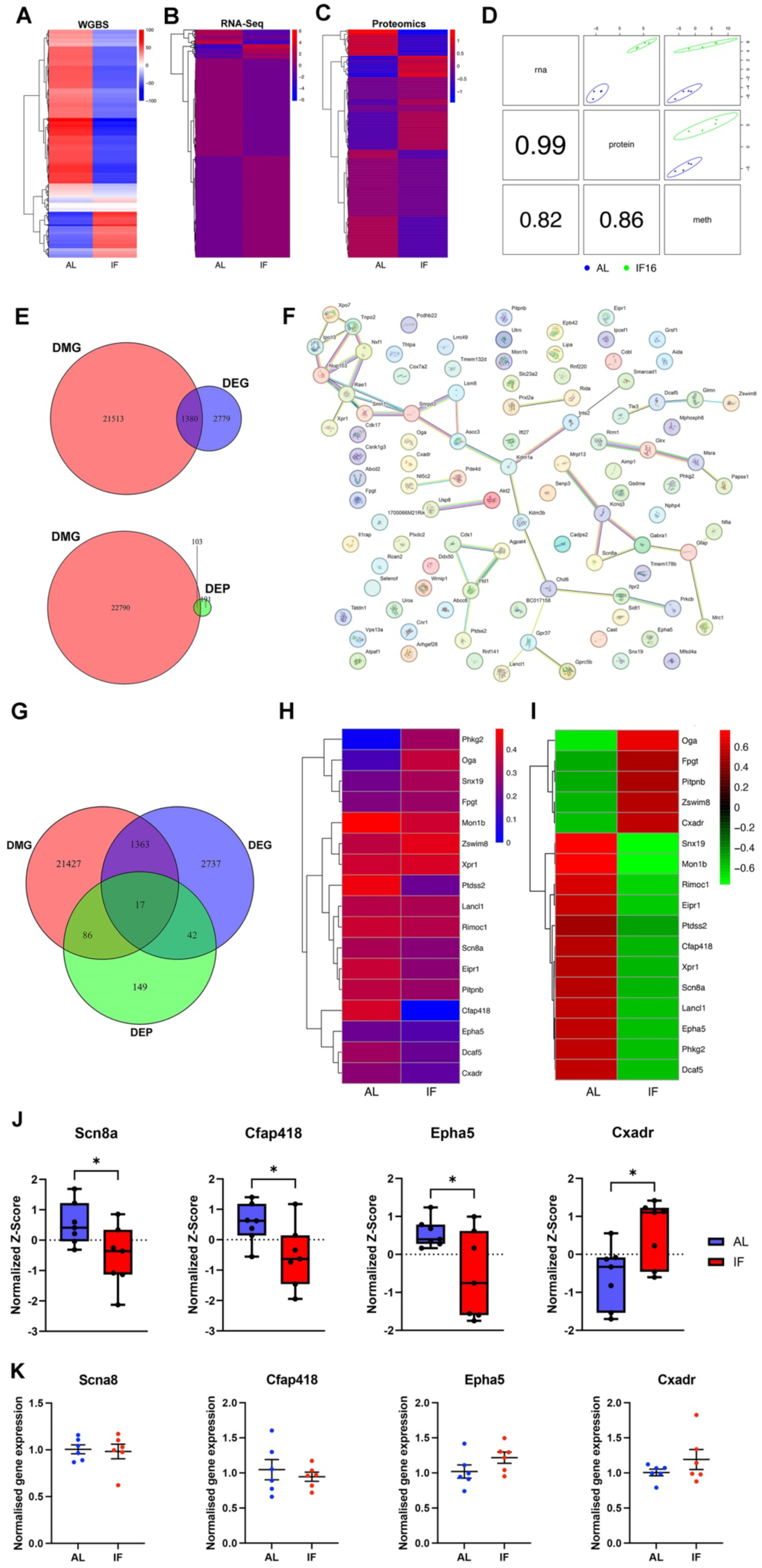
Integrated multiomic analysis reveals coordinated epigenetic, transcriptomic, and proteomic remodeling in the cortex of intermittent fasting fathers. (A–C) Heatmaps showing differential molecular signatures between ad libitum-fed (AL) and intermittent fasting (IF) paternal cortical samples. (A) Whole-genome bisulfite sequencing (WGBS) heatmap demonstrating differentially methylated genes (DMGs). (B) RNA-sequencing heatmap showing differentially expressed genes (DEGs). (C) Proteomic heatmap showing differentially expressed proteins (DEPs). (D) Pairwise correlation analysis integrating methylomic, transcriptomic, and proteomic datasets. Strong concordance was observed across molecular layers, indicating coordinated cortical remodeling induced by IF. (E) Venn diagram analyses showing overlap between DMGs and DEGs (top), and overlap between DMGs and DEPs (bottom). (F) Protein–protein interaction (PPI) network analysis of integrated candidate molecules identified across multiomic datasets. Network analysis revealed interconnected molecular modules associated with neuronal signaling, synaptic organization, intracellular transport, ion regulation, metabolism, and stress-response pathways. (G) Three-way Venn diagram illustrating overlap among DMGs, DEGs, and DEPs. Seventeen molecules were commonly altered across all three molecular layers, indicating coordinated epigenetic-transcriptional-proteomic regulation induced by IF. (H) Heatmap showing integrated methylome-transcriptome candidate genes exhibiting coordinated molecular alterations between AL and IF cortical samples. Representative molecules included *Scn8a, Epha5, Ptpnb, Dcaf5, Rimoc1,* and *Cxadr*. (I) Heatmap showing integrated methylome-proteome candidate proteins displaying coordinated differential expression patterns between AL and IF cortical samples. Representative proteins included *Pitpnb, Rae1, Cxadr, Oga, Gfap, Gprc5b,* and *Scn8a*. (J) Boxplot validation of representative integrated candidate molecules including *Scn8a, Cfap418, Epha5,* and *Cxadr*. IF significantly reduced expression of *Scn8a, Cfap418,* and *Epha5*, while *Cxadr* expression was significantly elevated relative to AL controls. Data are presented as normalized Z-scores; *P < 0.05.

To identify overlapping molecular signatures across datasets, Venn diagram analyses were performed. Comparison between differentially methylated genes (DMGs, i.e., genes with promoters identified as DMRs) and differentially expressed genes (DEGs) identified 1,380 overlapping genes (**Fig. 2E, top**), while integration of DMGs with differentially expressed proteins (DEPs) revealed 103 shared candidates (**Fig. 2E, bottom**). Protein–protein interaction (PPI) network analysis of integrated candidates revealed highly interconnected molecular modules associated with neuronal signaling, synaptic organization, cellular metabolism, ion transport, stress-response pathways, and cytoskeletal regulation (**Fig. 2F**). Several central hub molecules emerged within the network, including *Scn8a, Epha5, Itpr2, Gabral, Plxdc2,* and *Cxadr*, indicating potential regulatory roles in IF-induced cortical adaptation. Further three-way integration across methylome, transcriptome, and proteome datasets identified 17 common molecules altered across all three molecular layers (**Fig. 2G**), suggesting the existence of tightly coordinated epigenetic-transcriptional-proteomic regulatory networks induced by IF. Among the 17 overlapping candidates identified across the methylome, transcriptome, and proteomic datasets, DNA methylation analysis revealed distinct methylation patterns between the IF and AL groups, with several genes showing marked differences in methylation levels (**Fig. 2H and SI Appendix, Fig. S6**). These candidates included genes involved in neuronal signaling, ion channel regulation, metabolic adaptation, and membrane organization, such as *Scn8a, Epha5, Ptpnb, Rimoc1, and Cxadr*. Protein Z-score analysis further revealed distinct protein expression patterns between IF and AL cortical samples (**Fig. 2I and SI Appendix, Fig. S7**). Several proteins, including Pitpnb, Cxadr, Oga, Gfap, Gprc5b, and Scn8a, exhibited pronounced differences in protein abundance between the two groups. IF significantly reduced the protein levels of Scn8a, Cfap418, and Epha5, whereas Cxadr was significantly increased relative to AL controls (**Fig. 2J**). To further assess these findings, we performed qPCR analysis of the corresponding four transcripts identified from the RNA-seq data. However, qPCR did not reveal significant differences in transcript abundance between the AL and IF groups (**Fig. 2K**), suggesting that the observed differences in these candidates may be more prominently reflected at the proteomic than the transcriptomic level. Furthermore, integrated multiomic analysis using mixOmics demonstrated clear segregation between AL and IF cortical samples across methylomic, transcriptomic, and proteomic datasets (**SI Appendix, Fig. S8**). Hierarchical clustering revealed coordinated molecular remodeling induced by IF, with distinct integrated signatures observed between dietary groups. Several neuronal, metabolic, and stress-response associated molecules contributed prominently to group separation, supporting strong concordance across DNA methylation, RNA expression, and protein abundance datasets.

### Offspring of intermittent fasting parents exhibit widespread cortical DNA methylation remodeling

To determine whether parental IF induces intergenerational epigenetic remodeling in offspring, WGBS was performed on cerebral cortex tissue collected from female and male offspring (F_1_) derived from AL and IF parental groups. Genome-wide analysis identified widespread differentially methylated regions (DMRs) across all autosomes and sex chromosomes in both female (**Fig. 3A**) and male (**Fig. 3B**) IF offspring compared to AL offspring. The distribution of DMRs differed between sexes, with female offspring of IF parents (F_1_(IFXIF)) compared to female offspring of AL parents (F_1_(ALXAL)) exhibiting a predominance of hypermethylated regions (**Fig. 3A**), whereas male offspring of IF parents compared to male offspring of AL parents displayed predominantly hypomethylated (**Fig. 3B**) DMRs, indicating sex-specific intergenerational epigenomic remodeling induced by parental IF. Global methylation distribution analysis further demonstrated significant alterations in methylation profiles between AL parents offspring and IF parents offspring in both female (**Fig. 3C**) and male (**Fig. 3D**) groups. Hierarchical clustering analysis differentially methylated genes revealed clear segregation of cortical methylation signatures according to parental dietary condition in female (**Fig. 3E**) and male offspring (**Fig. 3F**), demonstrating robust intergenerational epigenomic remodeling induced by parental IF exposure.

**Figure 3.**
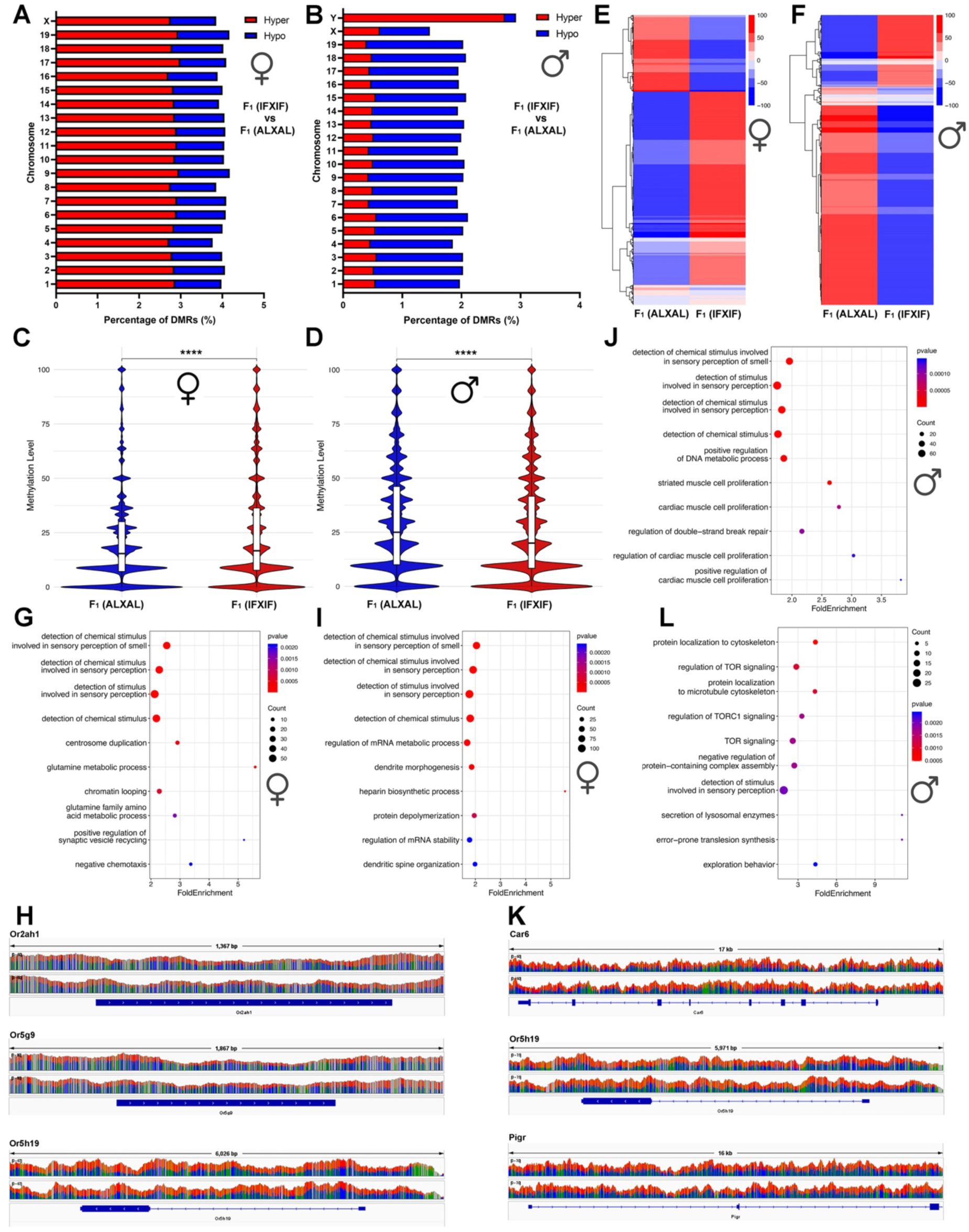
Offspring of intermittent fasting parents exhibit widespread cortical DNA methylation remodeling. (A,B) Chromosomal distribution of differentially methylated regions (DMRs) in female (A) and male (B) offspring derived from intermittent fasting (IF) parents relative to offspring derived from ad libitum (AL) parents. Female offspring displayed predominant hypermethylation-associated alterations, whereas male offspring exhibited a greater proportion of hypomethylated DMRs. (C,D) Violin plots showing global methylation level distributions between offspring derived from AL parents [F1(ALXAL)] and offspring derived from IF parents [F1(IFXIF)] in female (C) and male (D) offspring cortex samples. ****P < 0.0001. (E,F) Hierarchical clustering heatmaps demonstrating clear segregation of cortical methylation signatures according to parental dietary condition in female (E) and male (F) offspring. (G) Gene ontology (GO) enrichment analysis of hypomethylated genes in female offspring derived from IF parents, showing enrichment of sensory-associated pathways including olfactory signaling, sensory perception, glutamine metabolism, chromatin looping, and synaptic vesicle exocytosis. (H) Representative methylation tracks showing altered methylation patterns across sensory-associated loci including *Or2ah1, Or5g9,* and *Or5h19* in female offspring derived from IF parents relative to controls. (I) GO enrichment analysis of hypomethylated genes in male offspring derived from IF parents, showing enrichment of pathways associated with sensory perception, dendrite morphogenesis, mRNA metabolic processes, protein depolymerization, and dendritic spine organization. (J) GO enrichment analysis of hypermethylated genes in female offspring derived from IF parents, showing enrichment of sensory perception pathways together with regulation of DNA metabolic processes, cardiac muscle cell proliferation, and double-strand break repair pathways. (K) Representative methylation tracks showing altered methylation patterns across *Car6, Or5h19,* and *Pigr* loci in male offspring derived from IF parents relative to controls. (L) GO enrichment analysis of hypermethylated genes in male offspring derived from IF parents, showing enrichment of pathways associated with cytoskeletal organization, TOR/TORC1 signaling, lysosomal enzyme secretion, sensory perception, and exploratory behaviour.

GO enrichment analysis of differentially methylated genes revealed strong enrichment of sensory-associated pathways among differentially methylated genes in offspring derived from IF parents cerebral cortex compare to offspring derived from AL parents. In female offspring derived from IF parents, hypomethylated genes were enriched in pathways associated with sensory perception, detection of chemical stimulus involved in sensory perception of smell, centrosome duplication, glutamine metabolism, chromatin looping, and positive regulation of synaptic vesicle exocytosis (**Fig. 3G**). Representative methylation tracks further demonstrated altered methylation patterns across sensory-associated loci including Or2ah1, Or5g9, and Or5h19 in female offspring derived from IF parents (**Fig. 3H**). In female offspring derived from IF parents, hypermethylated genes were enriched in pathways associated with sensory perception, regulation of mRNA metabolic processes, dendrite morphogenesis, protein depolymerization, and dendritic spine organization (**Fig. 3I**). In male offspring derived from IF parents compare to male offspring derived from AL parents, hypomethylated genes were enriched in pathways associated with sensory perception, detection of chemical stimulus involved in sensory perception of smell, regulation of mRNA metabolic processes, dendrite morphogenesis, protein depolymerization, and dendritic spine organization (**Fig. 3J**). Representative methylation tracks further demonstrated reduced methylation across sensory-associated loci including Or5h19 in offspring derived from IF parents (**Fig. 3K**). In contrast, hypermethylated genes were enriched in pathways associated with protein localization to cytoskeleton, TOR signaling, regulation of TORC1 signaling, microtubule cytoskeleton organization, lysosomal enzyme secretion, and exploration behaviour (**Fig. 3L**). Additional representative methylation tracks demonstrated increased methylation across loci including Car6 and Pigr in IF-derived offspring relative to controls (**Fig. 3K**).

Heatmap analyses of differentially methylated genes further demonstrated extensive remodeling of sensory-associated gene families in offspring cortex (**SI Appendix, Fig. S9**). IF-derived female offspring exhibited prominent hypomethylation across multiple olfactory receptor and sensory signaling genes, including Gnat3, Grm6, Rtp2, Tas2r123, and Pigr (**SI Appendix, Fig. S9 A**), while hypermethylated genes included sensory-associated and neuronal signaling molecules such as Calca, Ryr2, Tac4, Cacna2d4, and Nrg1 (**SI Appendix, Fig. S9 B**). Similarly, IF-derived male offspring displayed extensive hypomethylation across olfactory receptor and sensory-associated genes including Nrg1, Piezo2, Car6, Grik2, and Pigr (**SI Appendix, Fig. S9 C**). In contrast, hypermethylated genes in male offspring included regulators associated with neuronal signaling, cytoskeletal organization, and metabolic pathways such as Hif1a, Gsk3b, Slc38a9, Capzb, and Ube3a (**SI Appendix, Fig. S9 D**). Collectively, these findings demonstrate that parental IF induces extensive intergenerational cortical DNA methylation remodeling in offspring, prominently affecting sensory-associated signaling pathways together with neuronal, cytoskeletal, and metabolic regulatory networks in both female and male descendants.

### Intergenerational multiomic remodeling reveals coordinated epigenetic, transcriptomic, and proteomic alterations in offspring cortex

To determine whether parental IF induces coordinated molecular remodeling across multiple regulatory layers in offspring, integrated analyses combining DNA methylation, transcriptomic, and proteomic datasets were performed using cerebral cortex tissue collected from female and male offspring. Hierarchical clustering of DMRs between male and female offspring derived from IF and AL parents showed clear segregation, revealing robust sex-specific epigenomic remodelling induced by parental IF exposure (**Fig. 4A**). Overlap analysis of hypomethylated and hypermethylated DMRs further identified distinct sex-dependent methylation signatures within the offspring cerebral cortex (**Fig. 4B**). IF-derived female offspring displayed predominant hypermethylation-associated alterations, whereas IF-derived male offspring exhibited a greater proportion of hypomethylated DMRs, indicating sexually dimorphic intergenerational epigenetic responses to parental IF. Proteomic analyses identified extensive remodeling of cerebral cortical protein expression profiles in both IF-derived female and IF-derived male offspring compared to AL-derived offspring. Volcano plot analyses demonstrated significant differential protein abundance between offspring derived from IF parents and offspring derived from AL parents in both females (**Fig. 4C**) and males (**Fig. 4D**). Pairwise integrative correlation analyses further revealed strong concordance between DMRs, transcriptomic and proteomic datasets in female (**Fig. 4E**) and male offspring (**Fig. 4F**), indicating coordinated intergenerational molecular remodeling across multiple regulatory layers.

**Figure 4.**
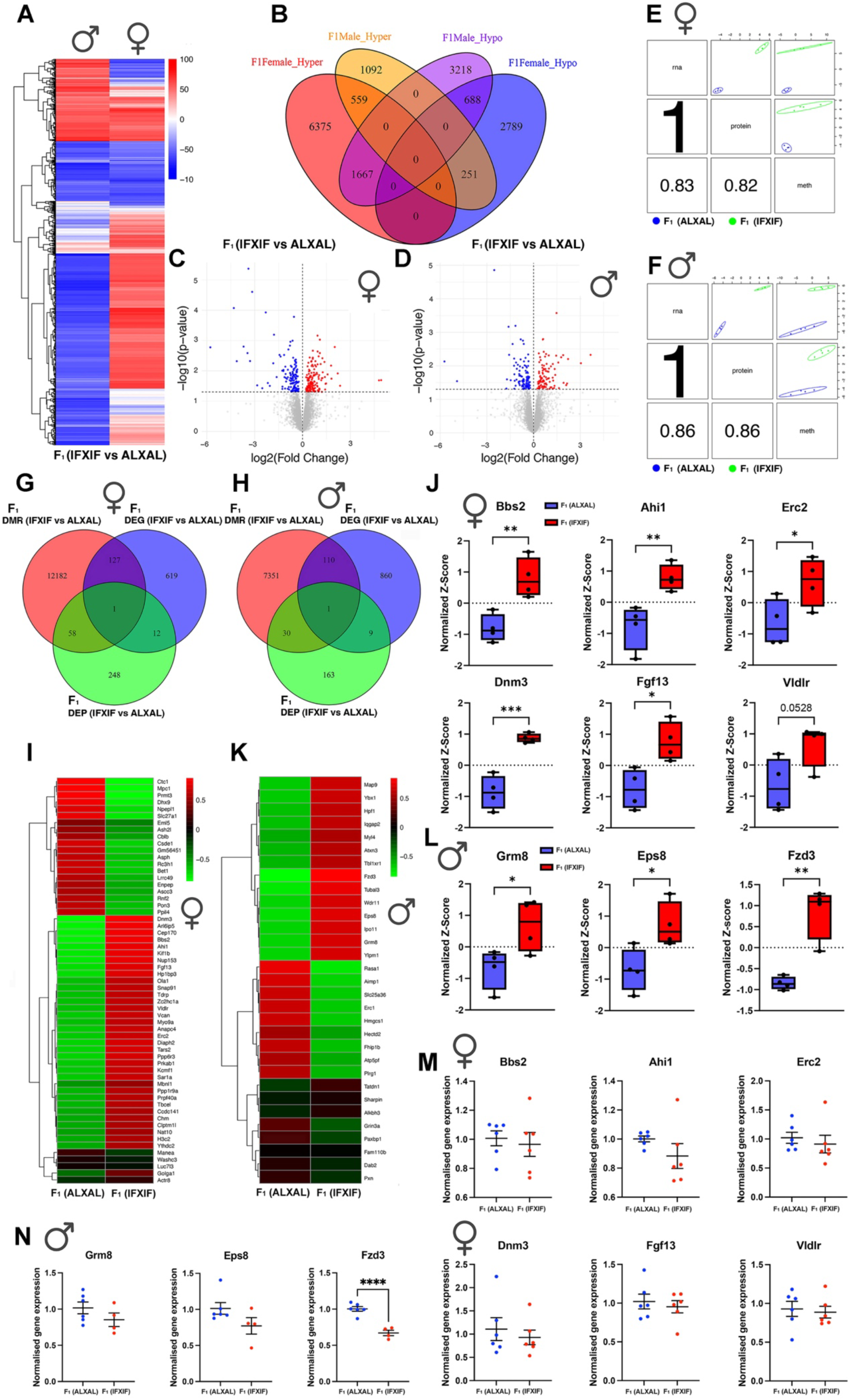
Intergenerational multiomic remodeling reveals coordinated epigenetic, transcriptomic, and proteomic alterations in offspring cortex. (A) Hierarchical clustering heatmap of differentially methylated regions (DMRs) demonstrating distinct sex-specific methylation signatures between offspring derived from ad libitum (AL) and intermittent fasting (IF) parents. (B) Venn diagram analysis showing overlap among hypermethylated and hypomethylated DMR-associated genes in female and male offspring derived from IF parents relative to offspring derived from AL parents. Female offspring exhibited predominant hypermethylation-associated alterations, whereas male offspring showed greater hypomethylation-associated signatures. (C,D) Volcano plot analyses of differentially expressed proteins (DEPs) in female (C) and male (D) offspring derived from IF parents compared with offspring derived from AL parents. Red dots indicate significantly upregulated proteins and blue dots indicate significantly downregulated proteins. (E,F) Pairwise integrative correlation analyses between methylomic, transcriptomic, and proteomic datasets in female (E) and male (F) offspring cortex samples, demonstrating coordinated molecular remodeling across regulatory layers. (G,H) Three-way Venn diagram analyses illustrating overlap among differentially methylated regions (DMRs), differentially expressed genes (DEGs), and differentially expressed proteins (DEPs) in female (G) and male (H) offspring derived from IF parents relative to controls. (I,K) Integrated DMR-DEP heatmaps showing coordinated protein expression signatures between offspring derived from AL and IF parents in female (I) and male (K) cortex samples. Female offspring displayed coordinated alterations in proteins associated with neuronal communication, synaptic organization, membrane signaling, and sensory-associated cortical pathways, including *Fgf13, Dnm3, Vldlr, Bbs2, Ahi1,* and *Erc2*. Male offspring exhibited differential expression of proteins associated with synaptic signaling, receptor trafficking, neuronal excitability, and sensory-associated neuronal pathways, including *Grm8, Eps8, Fzd3, Erc1, Adcy6,* and *Map9*. (J,L) Boxplot validation of representative candidate proteins in female (J) and male (L) offspring derived from IF parents relative to controls. Female offspring exhibited increased expression of *Bbs2, Ahi1, Erc2, Dnm3,* and *Fgf13*, while *Vldlr* showed a strong increasing trend. Male offspring demonstrated elevated expression of *Grm8, Eps8,* and *Fzd3*. Data are presented as normalized Z-scores; *P < 0.05, **P < 0.01, ***P < 0.001.

Three-way integration analyses identified overlapping candidate molecules shared across DMR, DEG, and DEP datasets in IF-derived female (**Fig. 4G**) and IF-derived male offspring (**Fig. 4H**) compared to AL-derived offspring. Female offspring exhibited one common overlapping candidate across all three molecular layers, together with substantial overlap between methylation and proteomic signatures (59 proteins) (**Fig. 4G**). Similarly, male offspring demonstrated coordinated overlap among methylation, transcriptomic, and proteomic datasets, supporting integrated intergenerational regulation across epigenetic, transcriptional, and protein levels. Overlap between methylation and proteomic signatures identified 31 proteins (**Fig. 4H**). Proteomic analyses further supported widespread cortical molecular remodeling in IF-derived offspring compared to AL-derived offspring (**SI Appendix, Fig. S10 A-D**). Volcano plot analyses identified extensive differential protein expression profiles in IF-derived female and male offspring compared to AL-derived offspring (**SI Appendix, Fig. S10 A, C**), while heatmap analyses demonstrated distinct clustering of proteomic signatures between offspring groups (**SI Appendix, Fig. S10 B, D**). In IF-derived female offspring, several proteins associated with sensory signaling, neuronal communication, membrane transport, and synaptic organization were differentially expressed, including Slc12a2, Ctnna3, Rasgrf2, Myo5c, Nlgn2, and Prph (**SI Appendix, Fig. S10 B**). In IF-derived male offspring, proteins linked to neurotransmission, receptor signaling, neuronal excitability, and synaptic organization, including Erc1, Adcy6, Grm8, Kcnv1, Pla2g6, and Sncg, exhibited marked differential expression between offspring groups (**SI Appendix, Fig. S10 D**), indicating substantial remodeling of sensory-associated neuronal pathways. RNA sequencing analyses similarly demonstrated widespread transcriptomic remodeling in offspring cortex (**SI Appendix, Fig. S11 A, B**).

Integrated DMR-DEP heatmap analyses revealed distinct integrated protein signatures between offspring derived from IF parents and controls (**Fig. 4I, K**). In IF-derived female offspring compared to AL-derived female offspring, coordinated alterations were observed in proteins associated with neuronal communication, synaptic organization, membrane signaling, cellular stress responses, and sensory-associated cortical pathways, including Fgf13, Dnm3, Vldlr, Bbs2, Ahi1, and Erc2 (**Fig. 4I**). Protein–protein interaction network analysis further demonstrated highly interconnected molecular modules involving RNA processing, intracellular trafficking, membrane organization, synaptic regulation, and neuronal signaling pathways (**SI Appendix, Fig. S12 A**). Several hub molecules, including Dhx9, Prpf40a, Snap91, Dnm3, and Bbs2, emerged as central regulators potentially contributing to sensory-associated cortical remodeling in female offspring. In IF-derived male offspring compared to AL-derived male, integrated analyses identified differential expression of proteins associated with synaptic signaling, receptor trafficking, neuronal excitability, cytoskeletal organization, and sensory-associated neuronal pathways, including Grm8, Eps8, Fzd3, Erc1, Adcy6, and Map9 (**Fig. 4K**). Corresponding protein–protein interaction analysis revealed interconnected signaling networks linked to glutamatergic signaling, intracellular transport, cytoskeletal dynamics, and synaptic plasticity (**SI Appendix, Fig. S12 B**). Central network molecules such as Eps8, Rasa1, Grm8, and Erc1 were prominently connected within these pathways, supporting coordinated intergenerational remodeling of sensory and neuronal signaling networks induced by parental IF exposure. Boxplot validation of proteomic data further confirmed significant alterations in representative candidate molecules associated with neuronal and sensory-associated signaling pathways. Female offspring exhibited significantly increased expression of Bbs2, Ahi1, Erc2, Dnm3, and Fgf13 in offspring derived from IF parents relative to controls, while Vldlr showed a strong increasing trend (**Fig. 4J**). In male offspring, proteins including Grm8, Eps8, and Fzd3 were significantly elevated in offspring derived from IF parents compared with controls (**Fig. 4L**), further supporting remodeling of glutamatergic signaling, synaptic plasticity, and neuronal connectivity pathways. To identify molecules shared between DMRs and DEPs, we next examined the corresponding transcripts and performed qPCR validation for the overlapping candidates. However, the qPCR results did not show significant differences between groups (**Fig. 4M and Fig. 4N**). This lack of concordance may reflect post-transcriptional and translational regulation, indicating that parental IF exposure may influence protein abundance and pathway activity without producing a detectable change in steady-state RNA levels. Together, these findings suggest that the observed epigenomic alterations may be functionally manifested primarily at the protein level. Global proteomic heatmap analysis identified extensive intergenerational remodeling of cortical protein expression profiles in offspring derived from IF parents relative to offspring derived from AL parents (**SI Appendix, Fig. S13**). In IF-derived female offspring (**SI Appendix, Fig. S13 A**), several differentially expressed proteins were associated with neuronal communication, sensory-associated signaling, membrane transport, and synaptic organization, including Slc12a2, Ctnna3, Rasgrf2, Myo5c, Nlgn2, Kirrel3, and Prph. Additional alterations in proteins such as Fgf13, Itgb8, and Nqo1 further suggest remodeling of neuronal excitability, intracellular signaling, and stress-response pathways. These coordinated changes indicate substantial adaptation of cortical sensory-processing and neuronal regulatory networks in female offspring following parental IF exposure. In IF-derived male offspring (**SI Appendix, Fig. S13 B**), parental IF induced differential expression of proteins associated with neurotransmission, glutamatergic signaling, receptor trafficking, neuronal excitability, and synaptic plasticity, including Grm8, Erc1, Adcy6, Kcnv1, Pla2g6, Sncg, and Map9. Several additional proteins linked to membrane dynamics, cytoskeletal regulation, oxidative stress responses, and intracellular transport, such as Dab2, Wdr11, H2bc6, and Ptp4a1, were also altered between offspring groups. Notably, proteins associated with sensory-associated neuronal signaling and synaptic communication pathways were prominently represented among the top differentially expressed proteins, suggesting persistent intergenerational remodeling of cortical sensory and neuronal network organization induced by parental IF. Collectively, these findings demonstrate that parental intermittent fasting induces coordinated intergenerational remodeling across epigenetic, transcriptomic, and proteomic layers within the offspring cerebral cortex, prominently affecting sensory-associated signaling pathways together with synaptic organization, neuronal communication, cytoskeletal dynamics, and stress-response regulatory networks in a sex-dependent manner.

### Integrated intergenerational analyses identify conserved sensory-associated molecular signatures across paternal and offspring generations

To determine whether IF-induced molecular alterations were conserved across generations, integrated analyses were performed comparing paternal (F_0_) cortical datasets with female and male offspring (F_1_) multiomic profiles. Venn diagram analysis of differentially methylated regions (DMRs) identified substantial overlap between fathers and offspring groups (**Fig. 5A**). A total of 2,724 shared DMR-associated genes were common across IF F_0_ fathers, IF-derived male offspring, and IF-derived female offspring, indicating persistent intergenerational epigenetic remodeling induced by parental IF exposure. Heatmap analysis of top overlapping DMR-associated genes demonstrated distinct intergenerational methylation patterns between offspring sexes (**Fig. 5B**). Male offspring exhibited methylation signatures that largely mirrored the hypomethylation patterns observed in IF fathers, indicating conserved intergenerational epigenetic inheritance across generations. In contrast, female offspring displayed an opposing methylation profile, with many shared loci exhibiting hypermethylation relative to the paternal IF signatures. Representative genes showing these sex-dependent methylation patterns included Dop1a, Sorbs2, Cphx1, and Smok2b, suggesting that parental IF induces sexually dimorphic epigenetic remodeling within neuronal and sensory-associated cortical pathways.

**Figure 5.**
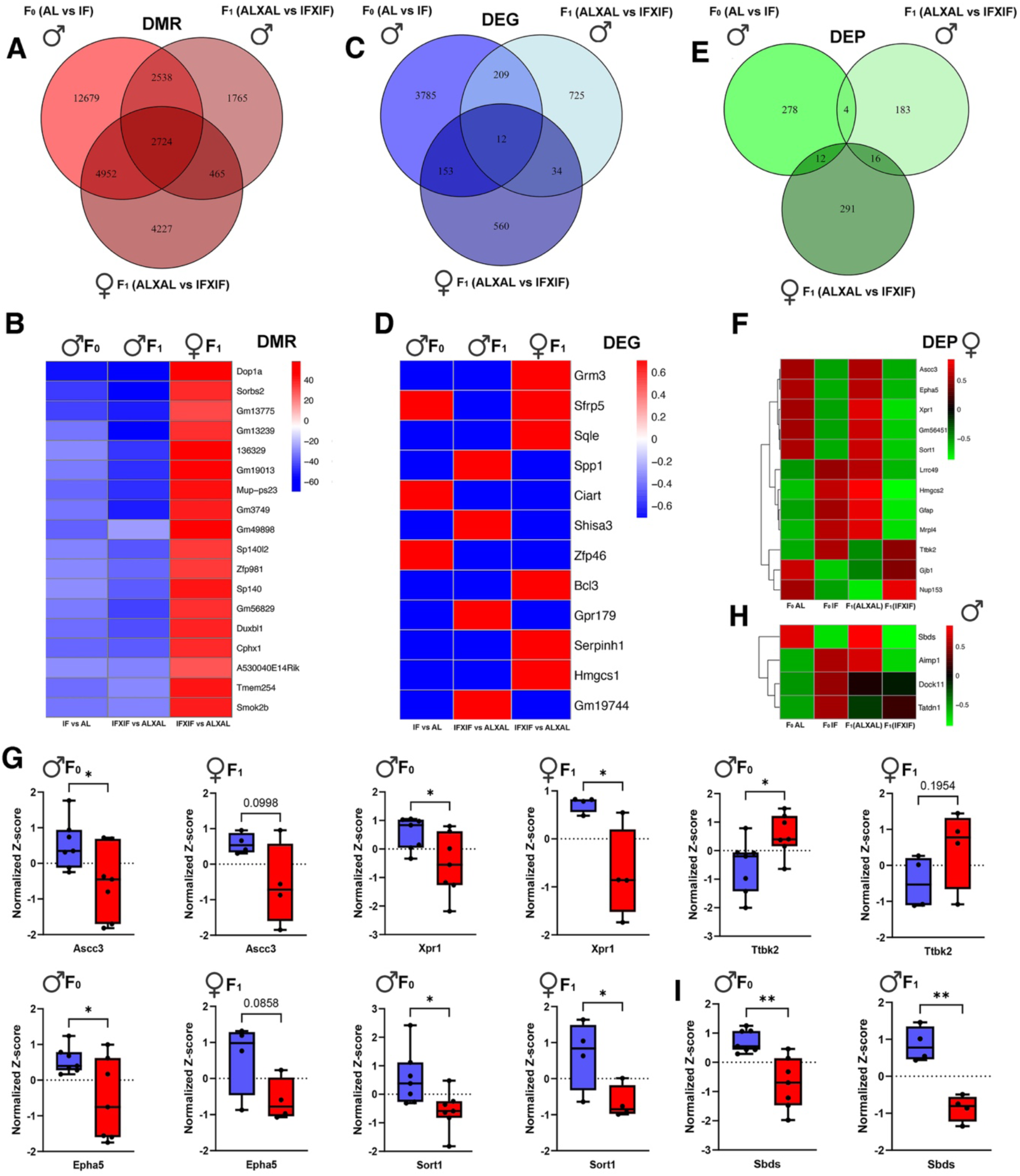
Integrated intergenerational analyses identify conserved sensory-associated molecular signatures across paternal and offspring generations. (A) Venn diagram analysis showing overlap of differentially methylated region (DMR)-associated genes between intermittent fasting (IF) fathers (F0), IF-derived male offspring, and IF-derived female offspring relative to corresponding ad libitum (AL) controls. A total of 2,724 shared DMR-associated genes were common across all groups. (B) Heatmap of representative overlapping DMR-associated genes demonstrating sex-dependent intergenerational methylation patterns. Male offspring exhibited methylation signatures that largely mirrored the hypomethylation patterns observed in IF fathers, whereas female offspring displayed opposing hypermethylation signatures. Representative genes included *Dop1a, Sorbs2, Cphx1,* and *Smok2b*. (C) Venn diagram analysis showing overlap of differentially expressed genes (DEGs) between IF fathers, IF-derived male offspring, and IF-derived female offspring. Twelve DEGs were shared across all three groups. (D) Heatmap of representative overlapping DEGs demonstrating sex-specific intergenerational transcriptional remodeling. Male offspring generally exhibited expression profiles similar to IF fathers, whereas female offspring displayed opposing expression patterns. Representative genes included *Grm3, Sfrp5, Sqle, Spp1, Ciart, Shisa3, Zfp46,* and *Serpinh1*. (E) Venn diagram analysis showing overlap of differentially expressed proteins (DEPs) between IF fathers, IF-derived male offspring, and IF-derived female offspring. Shared proteins were identified across paternal and offspring generations. (F,H) Heatmaps showing representative overlapping DEPs between IF fathers and IF-derived female offspring (F), and between IF fathers and IF-derived male offspring (H). Shared proteins included molecules associated with neuronal signaling, sensory-associated pathways, intracellular trafficking, cytoskeletal organization, and synaptic regulation, including *Ascc3, Epha5, Xpr1, Sort1, Ttbk2,* and *Sbds*. (G) Boxplot validation of representative conserved proteins between IF fathers and IF-derived female offspring, including *Ascc3, Epha5, Xpr1, Sort1,* and *Ttbk2*. (I) Boxplot validation showing reduced *Sbds* expression in both IF fathers and IF-derived male offspring relative to corresponding AL controls. Data are presented as normalized Z-scores; *P < 0.05, **P < 0.01.

Integrated transcriptomic analysis similarly revealed overlapping DEGs between IF fathers and IF-derived offspring compared to AL-derived offspring (**Fig. 5C**). Twelve genes were commonly altered across IF fathers, If-derived male offspring, and female offspring, while additional shared transcriptional signatures were identified between paternal and offspring datasets. Heatmap analysis demonstrated coordinated expression changes in genes associated with synaptic regulation, circadian signaling, metabolism, and neuronal communication, including *Grm3, Sfrp5, Sqle, Spp1, Ciart,* and *Shisa3* (**Fig. 5D**). Notably, transcriptomic heatmap analysis revealed distinct sex-dependent expression patterns across generations (**Fig. 5D**). Male offspring generally exhibited expression profiles that paralleled those observed in F_0_ fathers, whereas female offspring frequently displayed opposing transcriptional signatures. Representative genes demonstrating these sexually dimorphic intergenerational expression patterns included Grm3, Sfrp5, Sqle, Spp1, Ciart, Shisa3, Zfp46, and Serpinh1. Several of these genes are associated with glutamatergic signaling, synaptic organization, circadian regulation, extracellular signaling, and neuronal communication, suggesting that parental IF induces sex-specific transcriptional remodeling within sensory-associated and neural regulatory pathways across generations.

Proteomic integration analyses further identified shared DEPs between fathers and IF-derived offspring compared to AL-derived offspring (**Fig. 5E**). Several proteins were commonly altered across IF fathers and IF-derived female offspring, indicating persistent intergenerational remodeling at the protein level. Heatmap analyses revealed coordinated changes in proteins associated with neuronal signaling, membrane trafficking, cytoskeletal organization, and sensory-associated pathways (**Fig. 5F, H**). Shared candidate proteins between IF fathers and IF-derived offspring included *Ascc3, Epha5, Xpr1, Sort1, Sbds* and *Ttbk2.* Many of these proteins are functionally linked to neuronal connectivity, intracellular transport, synaptic regulation, and neurodevelopmental signaling pathways. Normalized Z-score analyses of proteomic further confirmed conserved intergenerational protein alterations. In both fathers and female offspring, proteins including *Ascc3, Epha5, Xpr1,* and *Sort1* exhibited consistent directional changes between IF and AL groups (**Fig. 5G**). Similarly, *Ttbk2* demonstrated coordinated alterations in IF fathers and IF-derived female offspring, while *Sbds* expression was significantly reduced in both IF fathers and IF-derived male offspring (**Fig. 5H, I**). Several of these molecules have established roles in neuronal maintenance, synaptic organization, intracellular trafficking, and sensory-associated cortical signaling. Collectively, these findings demonstrate that IF induces conserved intergenerational molecular signatures across epigenetic, transcriptomic, and proteomic layers in both fathers and offspring. Importantly, many of the shared molecules are associated with sensory-associated neuronal pathways, synaptic communication, intracellular transport, and cortical network regulation, supporting the existence of persistent multigenerational sensory-related molecular remodeling induced by parental IF exposure.

## Discussion

This study provides evidence that parental IF produces broad molecular remodeling in the cerebral cortex of both fathers and offspring, with a striking enrichment of sensory-associated pathways. In the F_0_ paternal brain, IF induced extensive changes in DNA methylation together with coordinated transcriptomic and proteomic remodeling. Across the cortex, the strongest signals repeatedly converged on genes and proteins involved in olfactory and taste perception, synaptic organization, glutamatergic signaling, cytoskeletal regulation, intracellular trafficking, and stress-response pathways. In the offspring generation, parental IF similarly induced widespread cortical remodeling across epigenetic, transcriptomic, and proteomic layers, with sex-dependent enrichment of sensory-associated, synaptic, and neuronal signaling pathways in both female and male offspring. These findings extend the biological impact of IF beyond classical metabolic adaptation and indicate that dietary timing can reshape neural molecular architecture in ways that are relevant to sensory processing and cortical function.

These findings are in line with prior studies showing that parental environmental interventions can be transmitted to offspring across generations. Paternal exercise has been shown to improve offspring brain physiology, adult neurogenesis, and cognition, with evidence that these benefits are inherited through germline-mediated epigenetic mechanisms [28]. Likewise, paternal stress has been shown to alter offspring HPA-axis regulation through sperm microRNA remodeling [29], and more recent work has reported reduced synaptic density and downregulation of learning-related genes in the offspring hippocampus and cortex [30, 31].

A major strength of this work is the integration of WGBS, RNA sequencing, and proteomics across paternal cerebral cortex and F_1_ offspring cerebral cortex. Rather than examining a single molecular layer in isolation, the present study shows that IF-associated changes are detectable across multiple levels of regulation and that many of these changes converge on the same functional themes. In fathers, the overlap between methylation, RNA, and protein data supports a coordinated molecular response rather than a set of unrelated changes. In offspring, the presence of intergenerational alterations in both female and male cortex suggests that parental IF can influence the neural epigenome and downstream molecular programs in the next generation, even in the absence of direct fasting exposure. These observations are also aligned with maternal programming studies showing that periconception undernutrition induces persistent, tissue-specific epigenetic changes in the offspring hypothalamus [32], while maternal high-fat diet is associated with long-term offspring neurodevelopmental disruption and adverse brain and behavioral consequences [33]. Together, these studies support the broader principle that parental nutritional environment can imprint lasting molecular and neural phenotypes in descendants.

The sensory component of these data is of particular interest. Olfactory and taste receptor-related genes, together with sensory transduction pathways and glutamatergic synaptic modules, were repeatedly enriched across the paternal and offspring datasets. This suggests that IF may alter the molecular basis of sensory information processing, which could have consequences for feeding behavior, environmental perception, and neural adaptation. Sensory systems are tightly linked to metabolic state, and these findings raise the possibility that IF may tune sensory circuitry to match energy availability and long-term physiological state [31, 34–37]. The concurrent enrichment of synaptic plasticity, neuronal communication, and cytoskeletal pathways further suggests that IF may influence not only sensory receptor networks but also the broader cortical machinery that interprets and integrates sensory input. Importantly, parental sensory experience itself can be inherited: olfactory fear conditioning in parents altered offspring behavioral sensitivity and olfactory neuroanatomy, with accompanying sperm DNA methylation changes at olfactory receptor loci [31]. This prior work provides a particularly strong conceptual precedent for our finding that sensory-associated pathways are among the most consistently remodeled systems after parental IF.

The offspring data add an important intergenerational dimension. The fact that F_1_ cerebral cortex retained molecular signatures related to sensory and neuronal pathways indicates that parental IF can transmit durable information across generations. Notably, the offspring responses were sex dependent, with female and male offspring showing different methylation and expression patterns. This dimorphism is biologically meaningful and may reflect sex-specific epigenetic inheritance, hormonal influences, or differences in developmental sensitivity to ancestral dietary history. The opposite directionality seen in some shared loci between male and female offspring suggests that intergenerational transmission is not uniform, but instead is shaped by sex-specific regulatory programs. Taken together, these convergent studies support the view that preconception environmental state can be encoded in gametes and expressed as offspring brain phenotypes, providing a strong precedent for the intergenerational sensory remodeling observed here. A parsimonious interpretation is that parental IF alters germline and/or maternal physiological information, potentially through coordinated changes in DNA methylation, chromatin state, and small RNA content, which may contribute to the sex-dependent cortical remodeling observed in offspring [29, 31, 38, 39]. Together, these findings provide a framework for understanding IF as a regulator of neural plasticity at both the individual and intergenerational levels. They also suggest that sensory remodeling may be one of the earliest and most conserved neural responses to dietary timing interventions. Because sensory pathways sit at the interface between external environment and internal physiology, their reprogramming may have broad effects on behavior, metabolic regulation, and brain resilience.

## Limitations

While the present study provides a comprehensive multiomic framework for understanding the intergenerational effects of IF on the brain, several limitations should be considered. The strong correlations observed across DNA methylation, transcriptomic, and proteomic datasets demonstrate coordinated molecular remodeling; however, the current analyses do not establish direct causal relationships between these regulatory layers. Additional mechanistic studies will therefore be necessary to determine whether the identified epigenetic alterations directly drive downstream transcriptional and proteomic changes. The analyses were conducted using bulk cerebral cortex tissue, which contains heterogeneous neuronal and glial populations. Consequently, the contribution of specific cell types to the observed molecular signatures could not be resolved. Future investigations using single-cell multiomics, spatial transcriptomics, or cell-type-specific epigenetic profiling will be important to define the precise cortical populations most responsive to IF and intergenerational programming.

Although substantial intergenerational molecular remodeling was identified in offspring, the biological mechanisms responsible for transmission across generations remain unclear. The present study did not directly examine germline epigenetic modifications, sperm-derived RNAs, early embryonic programming, or maternal developmental influences that may contribute to inheritance of IF-associated molecular signatures. An additional limitation is that, although both paternal and maternal animals underwent IF before mating, the present study did not experimentally disentangle paternal from maternal contributions to the observed offspring phenotypes and molecular signatures. Because IF was administered to both parents in the IF breeding group, the relative contributions of paternal germline transmission, maternal physiology, the intrauterine environment, and maternal postnatal influences cannot be determined from the current experimental design. Future studies should separately manipulate paternal and maternal IF, including reciprocal mating designs (e.g., IF♂ × AL♀ and AL♂ × IF♀), to distinguish paternal- versus maternal-mediated effects and determine their relative contributions to intergenerational programming. Such studies will also be important for defining whether maternal IF contributes independently or interacts with paternal IF to shape offspring cortical epigenetic, transcriptomic, proteomic, and sensory phenotypes.

The current work also focused primarily on molecular profiling. While the data strongly implicate sensory-associated, synaptic, and neuronal signaling pathways, additional functional studies examining sensory behavior, electrophysiological activity, synaptic plasticity, and neural circuit dynamics will be necessary to determine the physiological significance of these molecular alterations. In addition, the study examined a single IF paradigm within a defined experimental timeframe. Whether similar intergenerational cortical remodeling occurs under alternative fasting regimens, feeding schedules, ages, or environmental conditions remains unknown. Clear sex-specific molecular responses were also identified across multiple datasets, yet the hormonal and developmental mechanisms underlying these sexually dimorphic effects require further investigation.

## Materials and Methods

### Animals, IF procedures, Mating and Offspring Production

All in vivo experimental procedures were approved by the La Trobe University Animal Ethics Committee (AEC21047) and were conducted in accordance with the Australian Code for the Care and Use of Animals for Scientific Purposes (8th edition) and the NIH Guide for the Care and Use of Laboratory Animals. Every effort was made to minimize animal suffering and to reduce the number of animals used. All experiments were performed in accordance with the ARRIVE guidelines. C57BL/6 male and female mice were purchased at 4 weeks of age from ARC (Australia) and housed in the animal facility at La Trobe University. Mice were maintained on a 12 h light:12 h dark cycle (07:00–19:00) and provided with a standard diet containing 20% crude protein, 8.5% crude fat, 3.2% crude fiber, 1.1% calcium, and 0.96% phosphorus (Barastoc WEHI mouse breeder cubes Irradiated; Ridley, Australia). Water was available ad libitum to all groups. At 6 weeks of age, male and female mice were randomly assigned to either the intermittent fasting (IF; n = 64) group or the ad libitum feeding (AL; n = 64) control group. Mice in the IF group were subjected to a daily 16 h fasting regimen for 4 months (16:00–08:00), whereas AL mice had continuous access to food. Body weight was recorded weekly, and blood glucose and ketone concentrations were measured 6 h after fasting using the FreeStyle Optium Neo system with FreeStyle Optium blood glucose and ketone test strips (Abbott Laboratories, Illinois, USA). To minimize circadian variation, blood measurements were performed at the same time of day across all groups. After 4 months of dietary intervention, animals were paired for mating. To minimize potential selection bias, mating pairs were assigned randomly and without consideration of body weight or glucose levels. Male and female mice were assigned to parental dietary groups (AL or IF), and mating was performed within the corresponding treatment groups (AL father X AL mother; IF father X IF mother). Offspring were obtained and, at weaning, separated into male and female groups, with 30–50 animals per group across four groups in total. Both male and female offspring were maintained on a normal diet for 4 months. Following mating and weaning of the offspring, all parental animals were euthanized by carbon dioxide (CO2) inhalation between 07:00 and 12:00 h. Mice were then perfused with cold PBS, and tissues were collected, snap-frozen in liquid nitrogen, and stored at −80 °C until further analysis. At 6 months of age, offspring mice were humanely euthanized by carbon dioxide inhalation. The brain was harvested, the cerebral cortex was dissected, snap-frozen in liquid nitrogen, and stored at −80 °C until further analysis.

### DNA Extraction

Total DNA was extracted from cerebral cortex tissue using the QIAamp DNA Mini Kit (QIAGEN, USA) according to the manufacturer’s instructions. Approximately 20 mg of frozen cerebral cortex tissue was used for each sample, yielding 50 µl of DNA solution. Quality control was performed by measuring DNA concentration and the A260/A280 ratio using 1.5 µl of each sample on a NanoDrop ND-1000 spectrophotometer (Thermo Fisher Scientific, USA). Samples with a DNA concentration >20 ng/µl and an A260/A280 ratio >1.8 were considered to have passed QC and were submitted to the Australian Genome Research Facility (AGRF, Melbourne, Australia) for further quality control and sequencing.

### DNA Methylation Data Analysis

Raw reads from WGBS result were trimmed using Trim Galore v0.4.5 with the following options: clip_r1: 10, clip_r2: 10, three_prime_clip_r1: 10, and three_prime_clip_r2: 10. This step removed adapter/primer sequences and low-quality fragments. Mapping and methylation calls were performed using the DRAGEN Methylation Pipeline v4.03 in single-pass mode against the mm39 genome reference (mm39_GRCm39_Ensembl) and the spike-in Lambda DNA (*NC_001416.1*). Alignments were generated using the directional methylation protocol, with duplication marking and removal enabled. Methylation calls were produced using the cytosine and mbias report options. Methylation level, differentially methylated regions (DMRs) and differentially methylated genes (DMGs) were identified using methylkit package (1.38.0) on R-studio. Heatmap of differentially methylated genes were conducted by the pheatmap R package (1.0.12) with the average methylation level of each gene. Venn diagrams of overlapping genes were conducted by the VennDiagram R package (1.7.3). Gene Ontology (GO) enrichment analysis was performed using the ClusterProfiler R package (4.17.0).

### RNA Extraction

Total RNA was extracted from cortex sample using TRIzolTM RNA Isolation Reagents (Thermo Scientific, USA) following manufacturer’s instructions. 50 mg tissue was cut for each sample. RNA yield was determined by the NanoDrop One spectrophotometer (Thermo Scientific, USA). Extracted RNA samples were stored in −80°C and sent to Novogene (Singapore) for cDNA library preparation and RNA sequencing. Messenger RNA was purified from total RNA using poly-T oligo-attached magnetic beads. The library was checked with Qubit and real-time PCR for quantification and bioanalyzer for size distribution detection. Quantified libraries were pooled and sequenced on NovaSeq 6000 PE150.

### RNA Data Analysis

Raw data (raw reads) of fastq format were firstly processed through in-house perl scripts. Reference genome and gene model annotation files were downloaded from the genome website directly. Index of the reference genome was built using Hisat2 (2.0.5) and pairedend clean reads were aligned to the reference genome using Hisat2 (2.0.5). featureCounts (1.5.0-p3) was used to count the number of reads mapped to each gene. And then FPKM of each gene was calculated based on the length of the gene and reads count mapped to this gene. Differential expression analysis was conducted by DESeq2 R package (1.20.0). The P values were adjusted using the Benjamini & Hochberg method. Corrected P-value of 0.05 and absolute foldchange of 2 were set as the threshold for significantly differential expression.

### Protein extraction

Cortical samples from the parental group were homogenised and lysed in guanidine lysis buffer (6 M guanidine, 100 mM Tris, 10 mM TCEP, and 40 mM 2-chloroacetamide, pH 8.5), while cortical samples from the offspring group were lysed in SDS lysis buffer (4% SDS, 1 mM DTT, and 100 mM Tris-HCl, pH 7.6). Approximately 30–50 mg of cortical tissue was dissected, minced, and homogenised in 250 µL of the respective lysis buffer using a handheld homogeniser. Samples were then sonicated at 40% power for 15 cycles (15s on and 15s off), followed by incubation at 95°C for 5 min. The lysates were subsequently centrifuged at 20,000 × g for 30 min at 4°C, and the supernatants were collected. Protein concentrations were determined using the bicinchoninic acid (BCA) assay (Thermo Scientific, USA) according to the manufacturer’s instructions, with absorbance measured using a SPECTROstar Nano microplate reader (BMG LABTECH, Mornington, Victoria, Australia). Samples were then diluted to a final protein concentration of 0.5 µg/µL using the respective lysis buffer and submitted to the Proteomics and Metabolomics Platform at La Trobe University for proteomic analysis.

### Proteomics Sample Preparation and LC-MS/MS Analysis

Tissue lysates were processed using a modified SP3 (single-pot, solid-phase-enhanced sample preparation) approach optimized for efficient protein binding and digestion. Following reduction and alkylation, proteins were captured on carboxylate-coated magnetic beads in 50% ethanol, incubated at room temperature with agitation. After triple washes in 80% ethanol to remove contaminants, bound proteins were enzymatically digested overnight at 37 °C using trypsin in 20 mM ammonium bicarbonate at a 1:50 enzyme-to-protein ratio. Peptides were collected, acidified with 0.5% trifluoroacetic acid (TFA), purified via C18 StageTips, and dried under vacuum. For LC-MS/MS, peptides were reconstituted in 0.1% TFA and 5% acetonitrile (ACN), and 500 ng were loaded onto a C18 trap column, followed by separation on a 75 μm × 25 cm BEH analytical column with 1.7 μm particles. The gradient was run at 250 nL/min over 130 minutes using a Thermo Ultimate 3000 RSLCnano system, starting from 14% to 35% organic buffer (0.1% formic acid in 80% ACN), ramping to 50%, then to 95%. Mass spectra were acquired on a Tribrid Thermo Orbitrap Eclipse mass spectrometer, in DIA mode. Survey scans were collected at 60,000 resolution across 285–1200 m/z. DIA windows (4 m/z wide, staggered) covered the 285–1200 m/z range, acquired at 30,000 resolution using HCD fragmentation (30% normalized collision energy). Internal calibration was applied in realtime using EASY-IC.

### Protein Identification and Quantification

Raw DIA files were analyzed with DIA-NN 2.1.0 Academia, employing deep learning-based spectral prediction against the Mus musculus UniProt reference (September 2025 release). Searches specified trypsin/P digestion (up to two missed cleavages), with carbamidomethylation (C) as a fixed modification and methionine oxidation and Nterminal acetylation as variable. Peptides were filtered based on length (7-35 amino acids) and precursor mass range (285–1200 m/z). Protein inference and quantification adhered to a strict 1% FDR threshold at the protein level.

### Proteomics Data Analysis

Protein intensity data were log2-transformed and normalized using the normalizeMedianValues function from the limma R package to correct for inter-sample variation. Proteins with more than 50% missing values were discarded; the remaining missing entries were imputed using a left-shifted normal distribution approach, preserving variance structure. All downstream analyses were performed in the FlexStat web interface, which implements the empirical Bayes framework from limma for differential expression. Comparisons were made against the ad libitum (AL) control group, and significance was determined at an FDR-adjusted p < 0.05.

### Integrative Analysis

The integrative analysis between DNA methylation, RNA and proteomics data was conducted using mixOmics R package (6.36.0). Overlapping genes between each set were shown using Venndiagram R package (1.7.3). Heatmaps were generated by pheatmap R package (1.0.12).

### Quantitative Real-Time PCR Analysis

Total RNA was extracted using the RNeasy Lipid Tissue Mini Kit (Qiagen, Clayton, Australia), and cDNA was synthesised using the Tetro cDNA Synthesis Kit (Bioline, Boston, MA, USA) according to the manufacturer’s instructions. Quantitative real-time PCR (qPCR) was performed on a CFX96 Real-Time PCR Detection System (Bio-Rad, Hercules, CA, USA) using Luna Universal qPCR Master Mix (New England Biolabs, Ipswich, MA, USA). Each reaction was carried out in a final volume of 20 µL containing 0.25 µM of each primer and approximately 75 ng of cDNA. Amplification conditions consisted of an initial denaturation at 95°C for 1 min 15 s, followed by 45 cycles of 95°C for 1 min and 60°C for 30 s. Melt curve analysis was performed from 60°C to 95°C to confirm amplification specificity. All samples were analysed in technical triplicates, and a no-template control was included on each plate. Ct values were determined using CFX Manager software (Bio-Rad), and relative gene expression was calculated using the 2^-ΔΔCt method after normalisation to the reference genes Hprt1 and Rpl13a. Primer sequences are provided in Table below.

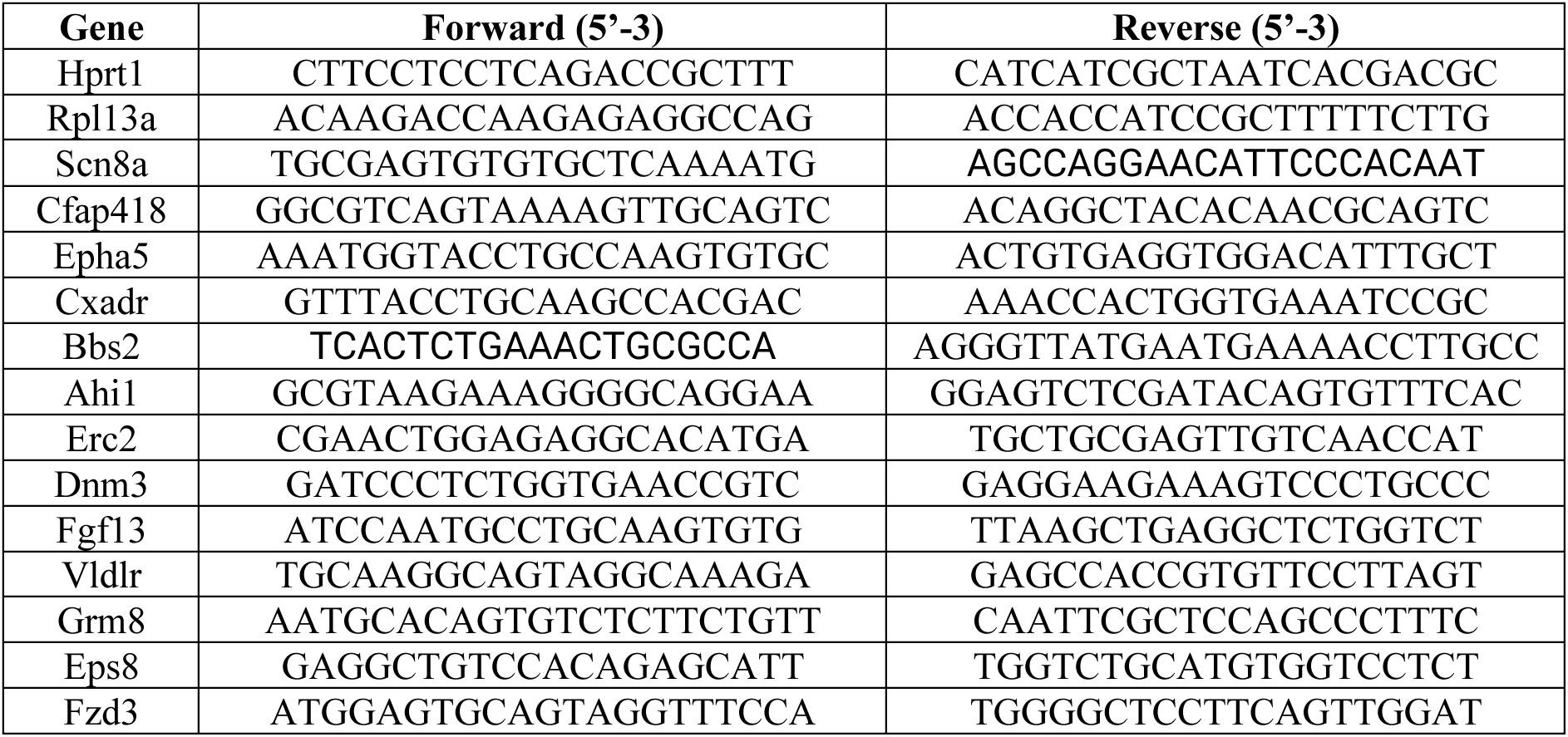

### Statistical Analysis

All the physiological parameter results were expressed as mean ± standard deviation (SD). GraphPad Prism (10.3.0) was used for statistical analysis. All the data were analysed with a T-test, one-way ANOVA, mixed-effects or two-way repeat-measurement ANOVA where required. The one-way ANOVA, mixed-effects or two-way repeated ANOVA were analysed with the following Tukey’s, Sidak’s or uncorrected Fisher’s LSD post hoc tests where required. P < 0.05 was considered statistically significant.

## Ethics approval

All *in vivo* experimental procedures were approved by La Trobe University (Ethics approval number: AEC21047) Animal Care and Use Committees and performed according to the guidelines set forth by the Australian Code for the Care and Use of Animals for Scientific Purposes (8th edition) and confirmed NIH Guide for the Care and Use of Laboratory Animals.

## Acknowledgments

This work was supported by La Trobe University (start-up grant to Thiruma V. Arumugam), the National Health and Medical Research Council of Australia (Grant Identification Number 2019100).

## Funding Information

This work was supported by the National Health and Medical Research Council of Australia (Grant Identification Number: 2019100) and Epigenes Pty. Ltd. Australia.

## Data availability

High-throughput RNA sequencing, whole-genome DNA methylation, and proteomic datasets generated in this study have been deposited in the appropriate public repositories, and the corresponding accession numbers will be provided upon availability.

## Notes

### Competing Interest Statement

The authors have declared no competing interest.

