## Supplementary Figures for "Integrated Multiomics Reveals Intergenerational Brain Sensory System Remodeling Induced by Parental Intermittent Fasting"

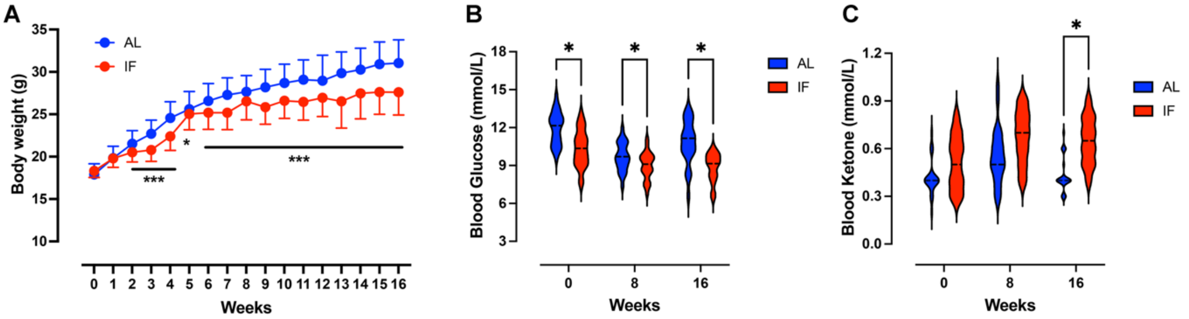


**Figure S1. Intermittent fasting induces physiological metabolic adaptation in paternal mice.** (A) Weekly body weight measurements of ad libitum-fed (AL) and intermittent fasting (IF) paternal mice over the 16-week dietary intervention period. IF mice exhibited significantly attenuated body weight gain compared with AL controls. (B) Violin plots showing blood glucose levels measured at 0, 8, and 16 weeks during the dietary intervention. IF mice displayed significantly reduced blood glucose levels relative to AL controls across time points. (C) Violin plots showing circulating blood ketone levels measured at 0, 8, and 16 weeks. IF mice exhibited significantly elevated ketone levels, indicating enhanced fasting-associated ketogenesis and metabolic switching. Data are presented as mean ± SD; *P < 0.05, ***P < 0.001.

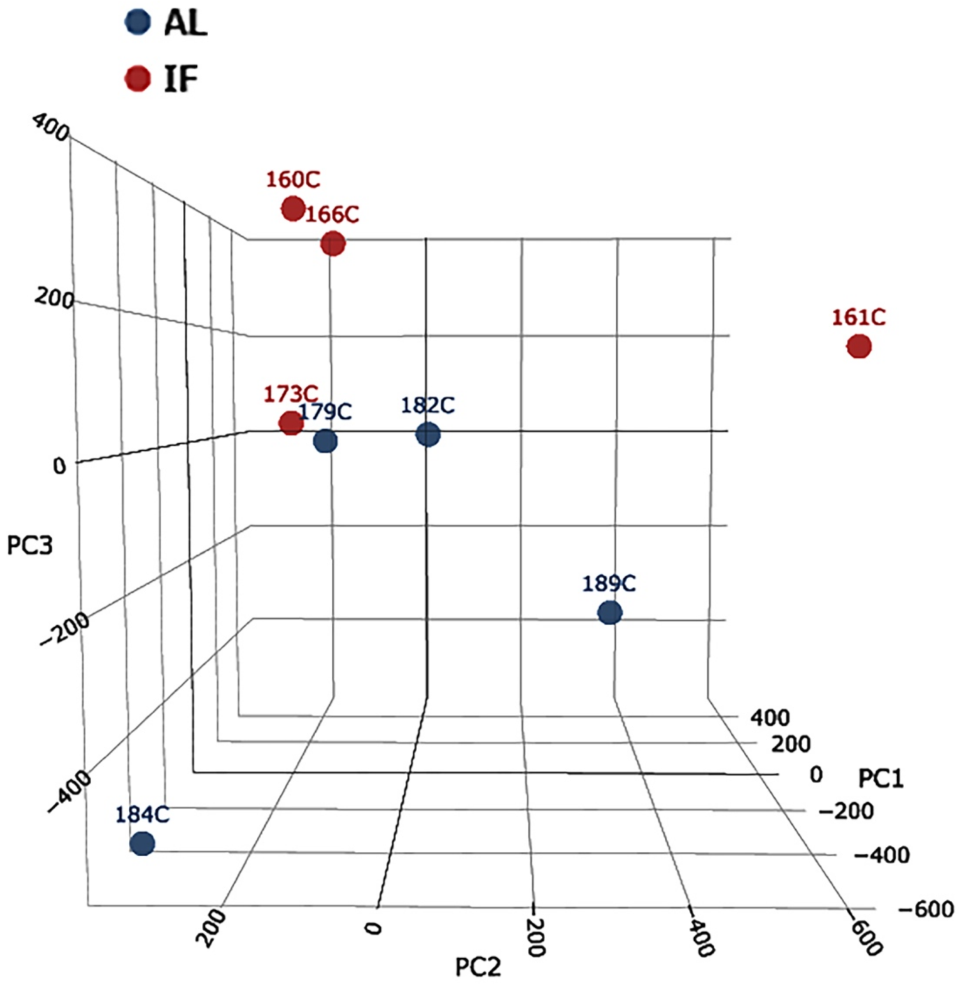


**Figure S2. Principal component analysis demonstrates distinct cortical methylation signatures induced by intermittent fasting in paternal mice.** Three-dimensional principal component analysis (PCA) plot of whole-genome bisulfite sequencing (WGBS) methylation profiles from ad libitum-fed (AL) and intermittent fasting (IF) paternal cortex samples. IF samples clustered separately from AL controls across principal component dimensions, indicating substantial fasting-induced divergence in global cortical DNA methylation signatures. Blue circles represent AL samples and red circles represent IF samples.

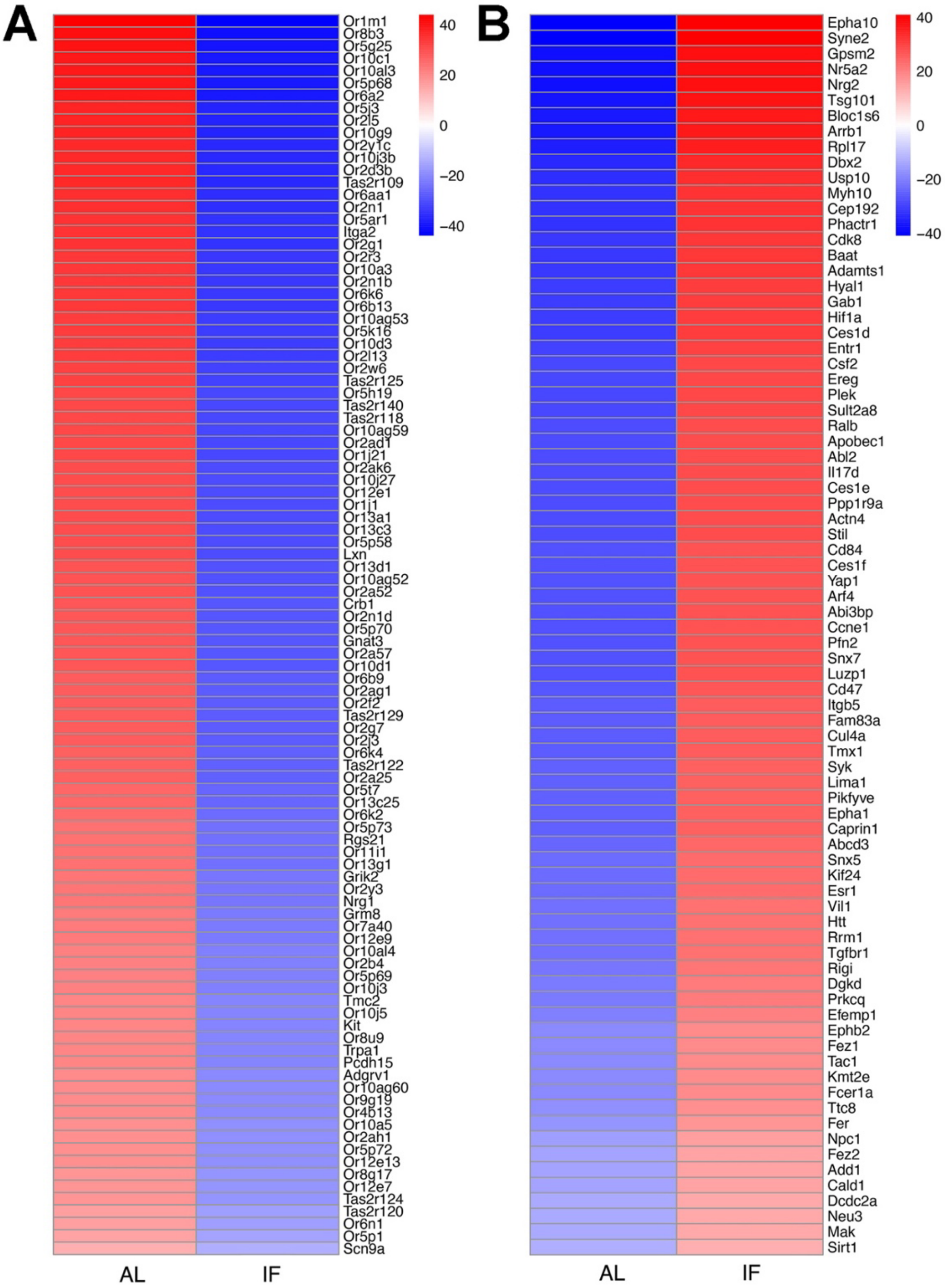


**Figure S3. Intermittent fasting induces extensive remodeling of sensory-associated and neuronal cortical methylation signatures in paternal mice.** (A) Heatmap of representative hypomethylated genes in intermittent fasting (IF) paternal cortex samples relative to ad libitum-fed (AL) controls. Numerous olfactory receptor and taste receptor-associated genes exhibited reduced methylation in IF cortex, including members of the *Olfr* and *Tas2r* gene families together with sensory signaling-related genes such as *Gnat3, Grik2,* and *Trpa1*. (B) Heatmap of representative hypermethylated genes in IF paternal cortex samples relative to AL controls. Hypermethylated genes included molecules associated with neuronal signaling, cellular organization, metabolic adaptation, and stress-response pathways, including *Epha10, Syn2, Nrg2, Hif1a,* and *Sirt1*. Color scale represents relative methylation levels.

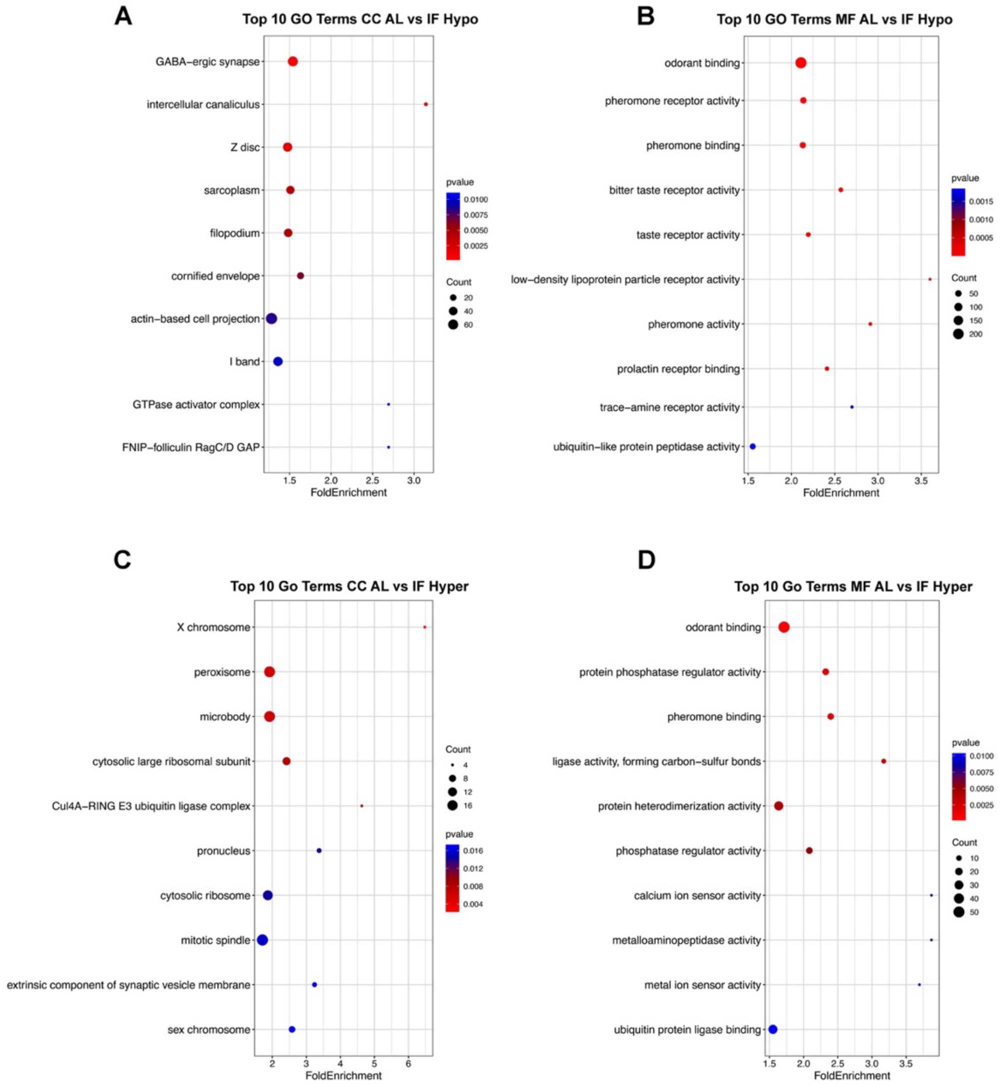


**Figure S4. Functional enrichment analyses reveal sensory-associated and neuronal pathway remodeling in paternal intermittent fasting cortex.** (A) Gene ontology (GO) cellular component enrichment analysis of hypomethylated genes in intermittent fasting (IF) paternal cortex samples relative to ad libitum-fed (AL) controls. Enriched pathways included GABAergic synapse, actin-based cell projections, filopodia, intercellular canaliculi, and cytoskeletal-associated structures. (B) GO molecular function enrichment analysis of hypomethylated genes showing strong enrichment of sensory-associated pathways including odorant binding, pheromone receptor activity, pheromone binding, bitter taste receptor activity, taste receptor activity, prolactin receptor binding, and trace-amine receptor activity. (C) GO cellular component enrichment analysis of hypermethylated genes in IF cortex samples. Enriched pathways included peroxisome, microbody, mitotic spindle, ribosomal complexes, and synaptic vesicle membrane-associated structures. (D) GO molecular function enrichment analysis of hypermethylated genes showing enrichment of odorant binding, pheromone binding, phosphatase regulator activity, calcium ion sensor activity, metallopeptidase activity, and ubiquitin protein ligase binding pathways. Bubble size represents gene count and color indicates adjusted P value.

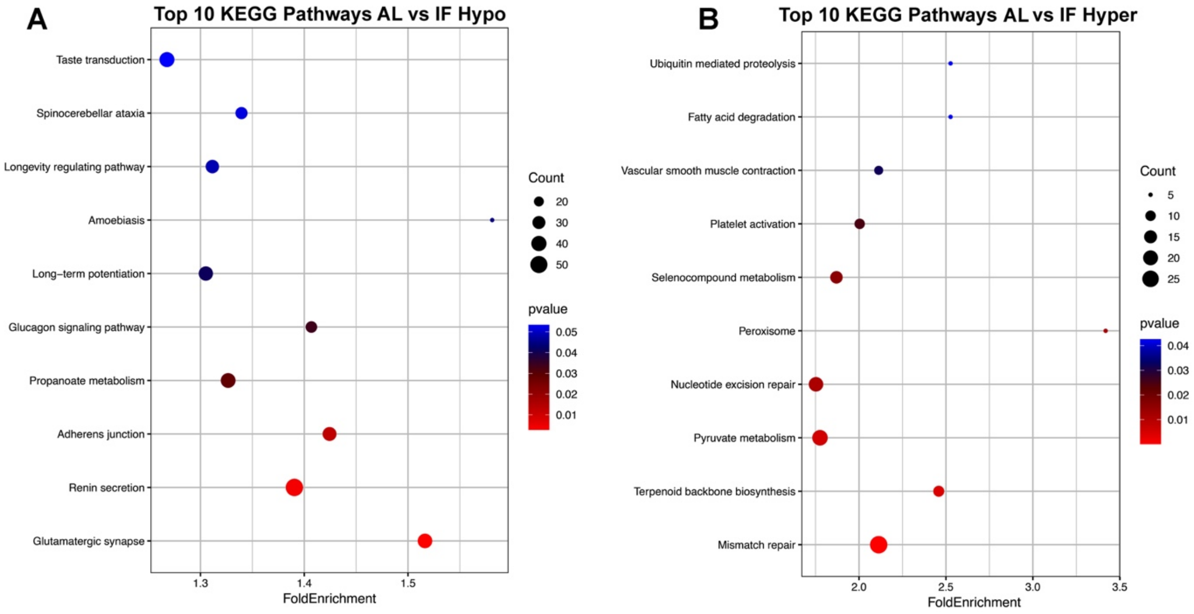


**Figure S5. KEGG pathway enrichment analysis reveals neuronal and metabolic pathway remodeling in paternal intermittent fasting cortex.** (A) KEGG pathway enrichment analysis of hypomethylated genes in intermittent fasting (IF) paternal cortex samples relative to ad libitum-fed (AL) controls. Enriched pathways included taste transduction, glutamatergic synapse, long-term potentiation, longevity regulating pathway, glucagon signaling pathway, renin secretion, and spinocerebellar ataxia pathways, indicating prominent remodeling of sensory-associated and neuronal signaling networks. (B) KEGG pathway enrichment analysis of hypermethylated genes in IF cortex samples. Enriched pathways included mismatch repair, nucleotide excision repair, pyruvate metabolism, fatty acid degradation, ubiquitin-mediated proteolysis, vascular smooth muscle contraction, terpenoid backbone biosynthesis, and peroxisome pathways, suggesting coordinated modulation of metabolic, stress-response, and cellular maintenance pathways. Bubble size represents gene count and color indicates adjusted P value.

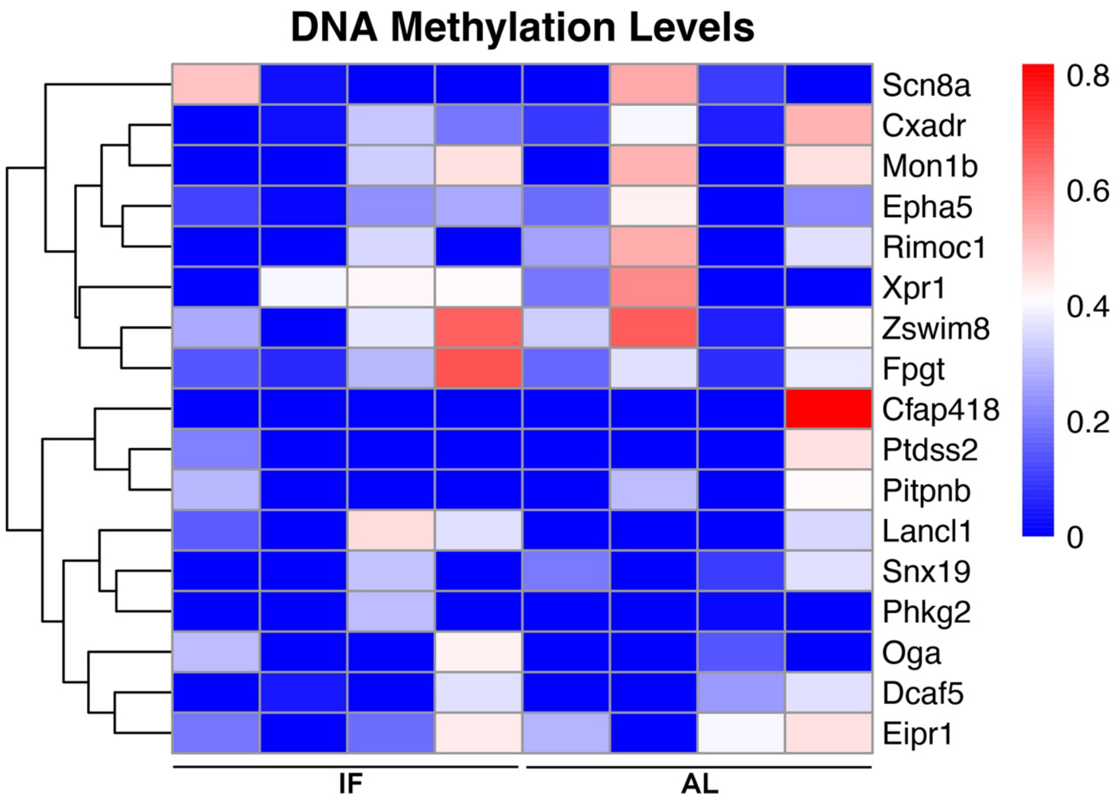


**Figure S6. Integrated methylome-transcriptome heatmap analysis reveals coordinated molecular remodeling in paternal intermittent fasting cortex.** Heatmap showing integrated candidate genes identified from overlapping methylome and transcriptome analyses in intermittent fasting (IF) paternal cortex samples relative to ad libitum-fed (AL) controls. Representative molecules exhibiting coordinated methylation and transcriptional alterations included *Scn8a, Cxadr, Mon1b, Epha5, Rimoc1, Xpr1, Zswim8, Cfap418, Pitpnb,* and *Dcaf5*. Many of these genes are associated with neuronal signaling, intracellular transport, membrane organization, synaptic regulation, and sensory-associated cortical pathways. Color scale represents relative DNA methylation levels.

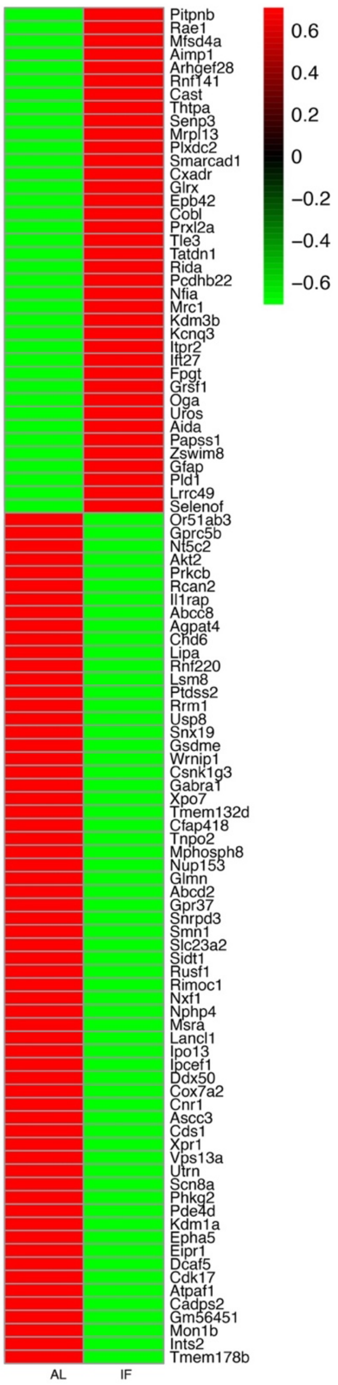


**Figure S7. Integrated methylome-proteome heatmap analysis reveals coordinated protein remodeling in paternal intermittent fasting cortex.** Heatmap showing integrated candidate proteins identified from overlapping methylome and proteomic analyses in intermittent fasting (IF) paternal cortex samples relative to ad libitum-fed (AL) controls. Representative proteins exhibiting coordinated differential expression patterns included *Pitpnb, Rae1, Cast, Senp3, Plxdc2, Cxadr, Itpr2, Kcnq3, Oga, Gfap, Gprc5b, Gabral, Scn8a, Epha5,* and *Cfap418*. Many of these proteins are associated with neuronal signaling, synaptic organization, intracellular transport, membrane dynamics, sensory-associated cortical pathways, and stress-response regulation. Color scale represents relative protein expression levels.

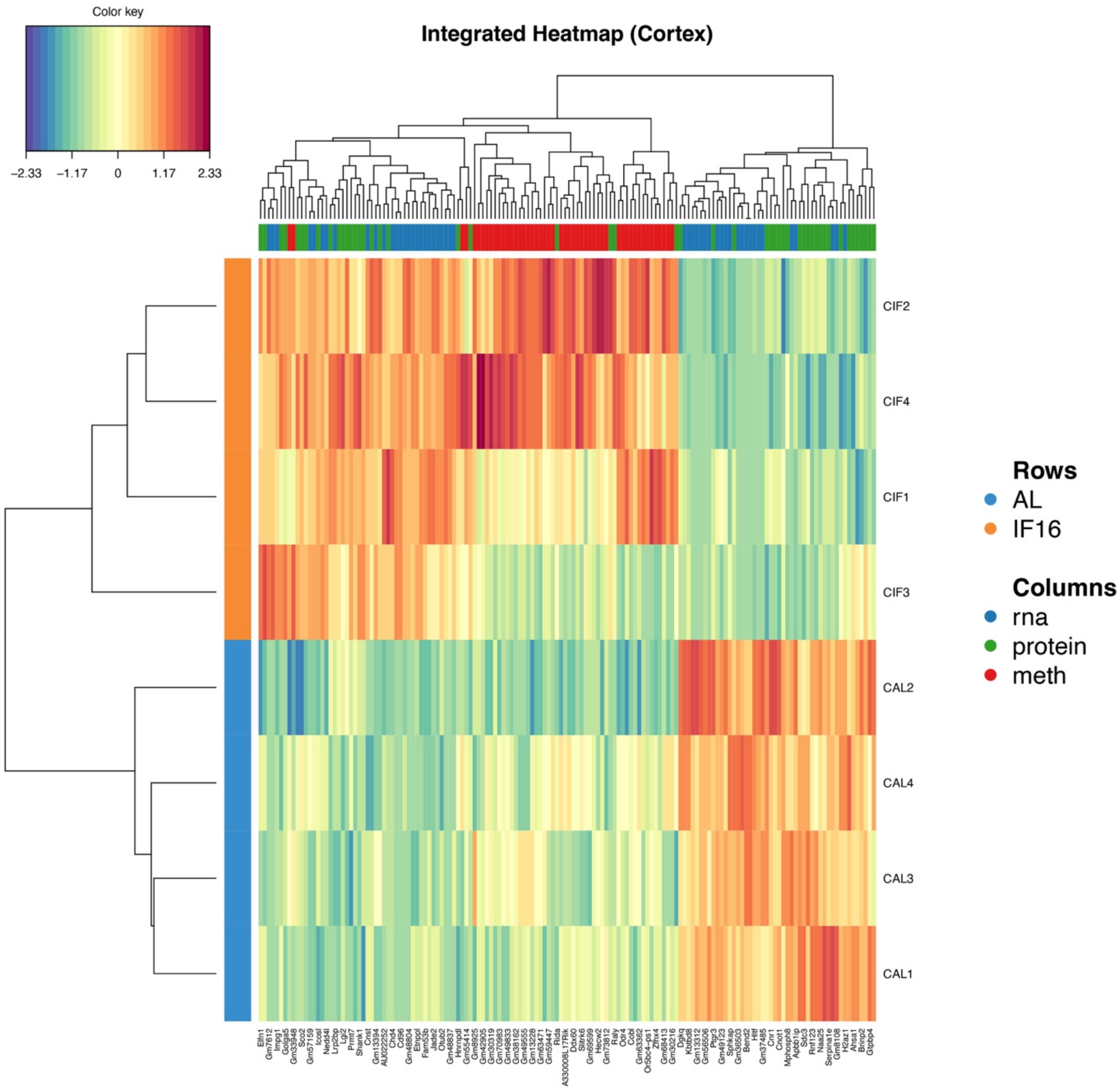


**Figure S8. mixOmics integration analysis reveals coordinated multiomic segregation between ad libitum and intermittent fasting paternal cortex samples.** Integrated multiomic heatmap generated using mixOmics analysis combining methylomic, transcriptomic, and proteomic datasets from ad libitum-fed (AL) and intermittent fasting (IF) paternal cortex samples. Hierarchical clustering demonstrated clear segregation between AL and IF groups across integrated molecular signatures, indicating coordinated remodeling across DNA methylation, RNA expression, and protein abundance datasets. Several neuronal, metabolic, and stress-response associated molecules contributed prominently to group separation, supporting strong concordance among epigenetic, transcriptomic, and proteomic regulatory layers. Rows represent individual cortical samples and columns represent integrated molecular features derived from RNA, protein, and methylation datasets.

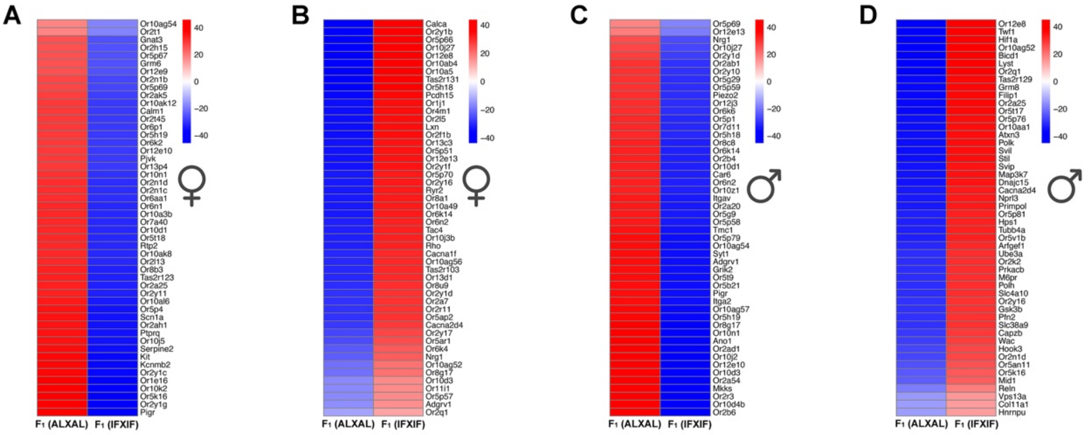


**Figure S9. Intergenerational remodeling of sensory-associated cortical methylation signatures in offspring derived from intermittent fasting parents.** (A) Heatmap of representative hypomethylated genes in female offspring derived from intermittent fasting (IF) parents relative to offspring derived from ad libitum (AL) parents. Prominent hypomethylation was observed across multiple olfactory receptor and sensory signaling-associated genes, including *Gnat3, Grm6, Rtp2, Tas2r123,* and *Pigr*. (B) Heatmap of representative hypermethylated genes in IF-derived female offspring. Hypermethylated genes included sensory-associated and neuronal signaling molecules such as *Calca, Ryr2, Tac4, Cacna2d4,* and *Nrg1*. (C) Heatmap of representative hypomethylated genes in male offspring derived from IF parents relative to controls. Extensive hypomethylation was observed across olfactory receptor and sensory-associated genes including *Nrg1, Piezo2, Car6, Grik2,* and *Pigr*. (D) Heatmap of representative hypermethylated genes in IF-derived male offspring. Hypermethylated genes included regulators associated with neuronal signaling, cytoskeletal organization, and metabolic pathways such as *Hif1a, Gsk3b, Slc38a9, Capzb,* and *Ube3a*. Color scale represents relative DNA methylation levels.

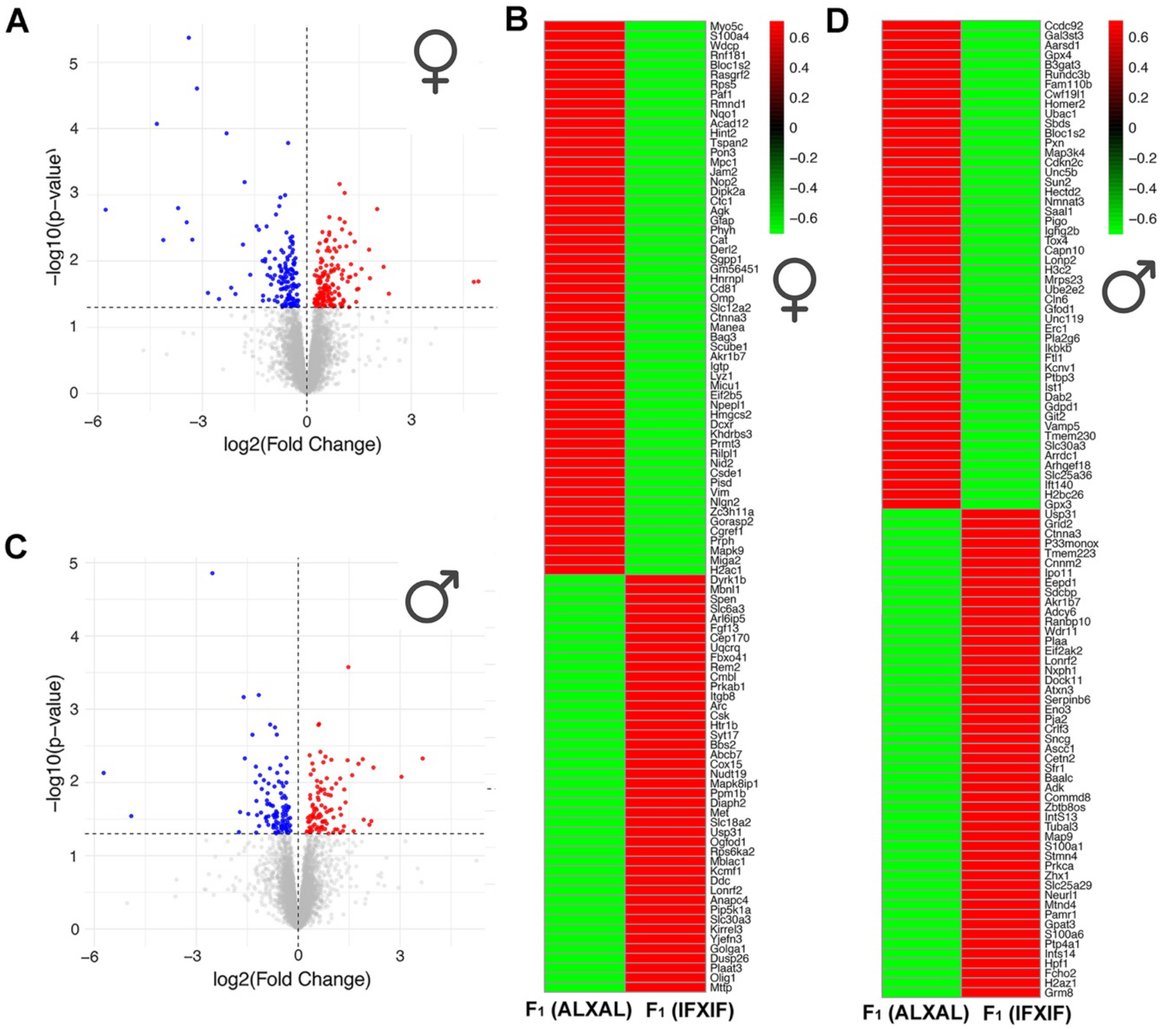


**Figure S10. Proteomic analyses reveal extensive sensory-associated cortical protein remodeling in offspring derived from intermittent fasting parents.** (A,C) Volcano plot analyses of differentially expressed proteins (DEPs) in female (A) and male (C) offspring derived from intermittent fasting (IF) parents relative to offspring derived from ad libitum (AL) parents. Red dots indicate significantly upregulated proteins and blue dots indicate significantly downregulated proteins. (B,D) Heatmaps showing representative differentially expressed proteins in female (B) and male (D) offspring cortex samples. In IF-derived female offspring, proteins associated with sensory signaling, neuronal communication, membrane transport, and synaptic organization were differentially expressed, including *Slc12a2, Ctnna3, Rasgrf2, Myo5c, Nlgn2,* and *Prph*. In IF-derived male offspring, proteins linked to neurotransmission, receptor signaling, neuronal excitability, and synaptic organization, including *Erc1, Adcy6, Grm8, Kcnv1, Pla2g6,* and *Sncg*, exhibited marked differential expression relative to controls. Color scale represents relative protein expression levels.

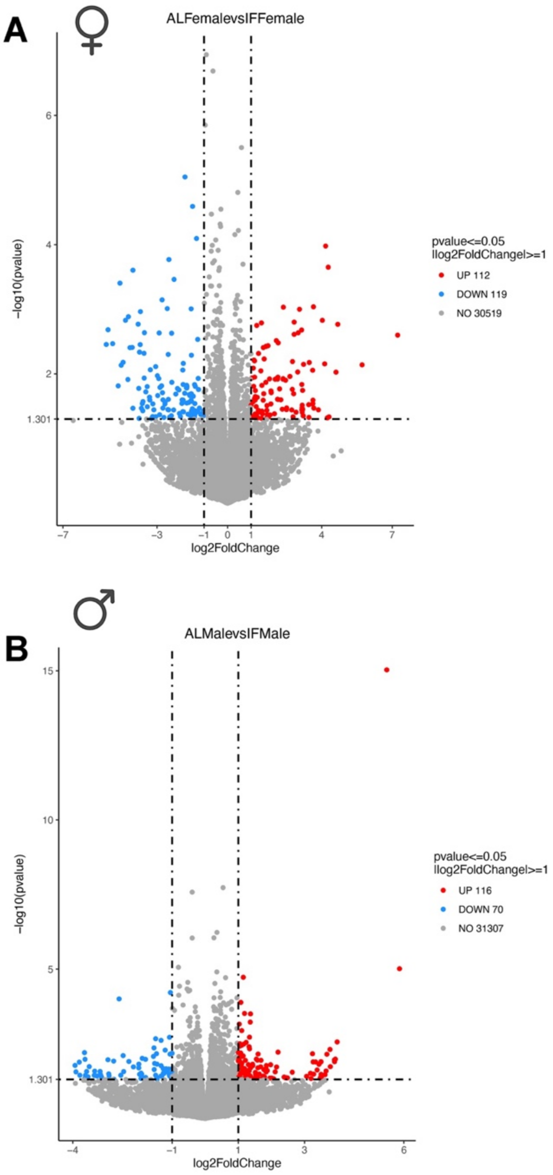


**Figure S11. RNA sequencing analyses reveal widespread transcriptomic remodeling in offspring derived from intermittent fasting parents.** (A,B) Volcano plot analyses of differentially expressed genes (DEGs) in female (A) and male (B) offspring derived from intermittent fasting (IF) parents relative to offspring derived from ad libitum (AL) parents. Red dots indicate significantly upregulated genes and blue dots indicate significantly downregulated genes. Female offspring exhibited 112 significantly upregulated and 119 significantly downregulated genes, whereas male offspring displayed 116 significantly upregulated and 70 significantly downregulated genes, indicating substantial intergenerational transcriptomic remodeling induced by parental IF exposure. Differentially expressed genes included molecules associated with neuronal signaling, sensory-associated pathways, synaptic organization, metabolism, and stress-response regulation.

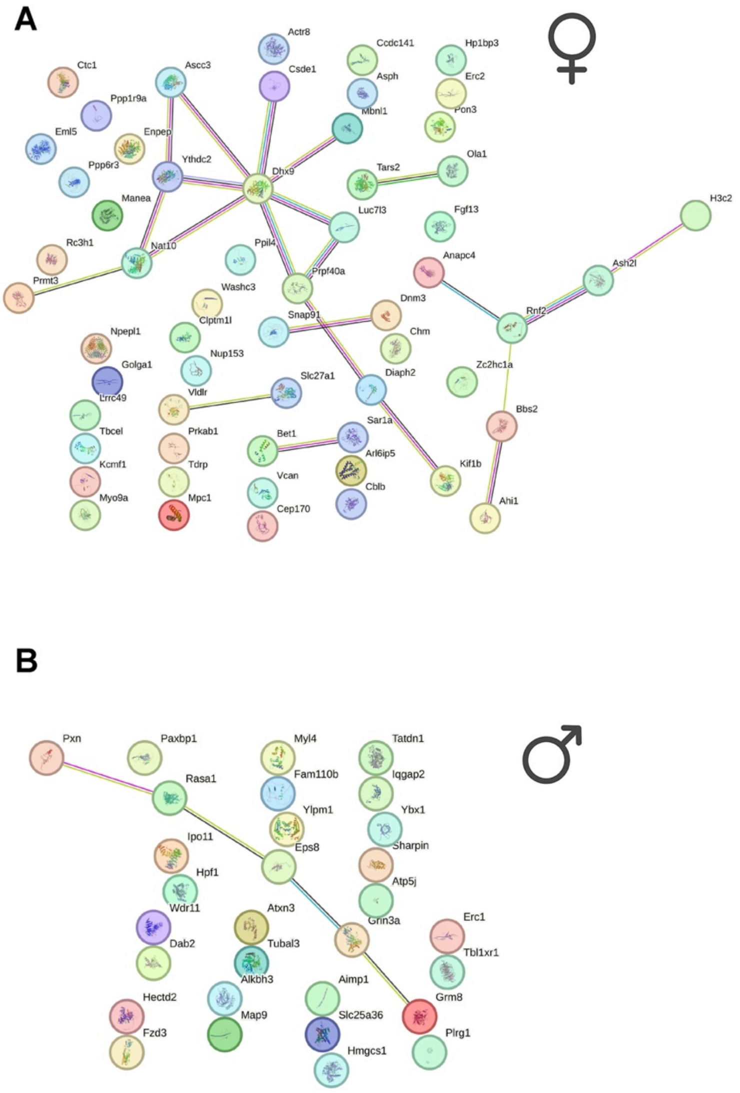


**Figure S12. Protein–protein interaction network analysis reveals interconnected sensory-associated and neuronal signaling modules in offspring derived from intermittent fasting parents.** (A) Protein–protein interaction (PPI) network analysis of integrated methylome-proteome candidate molecules in female offspring derived from intermittent fasting (IF) parents relative to offspring derived from ad libitum (AL) parents. Highly interconnected molecular modules were associated with RNA processing, intracellular trafficking, membrane organization, synaptic regulation, and neuronal signaling pathways. Central hub molecules included *Dhx9, Prpf40a, Snap91, Dnm3,* and *Bbs2*, suggesting coordinated remodeling of sensory-associated cortical pathways. (B) PPI network analysis of integrated methylome-proteome candidate molecules in male offspring derived from IF parents relative to controls. Interconnected molecular networks were associated with glutamatergic signaling, cytoskeletal dynamics, intracellular transport, neuronal excitability, and synaptic plasticity. Prominent hub molecules included *Eps8, Rasa1, Grm8,* and *Erc1*, supporting coordinated intergenerational remodeling of sensory-associated and neuronal signaling pathways induced by parental IF exposure.

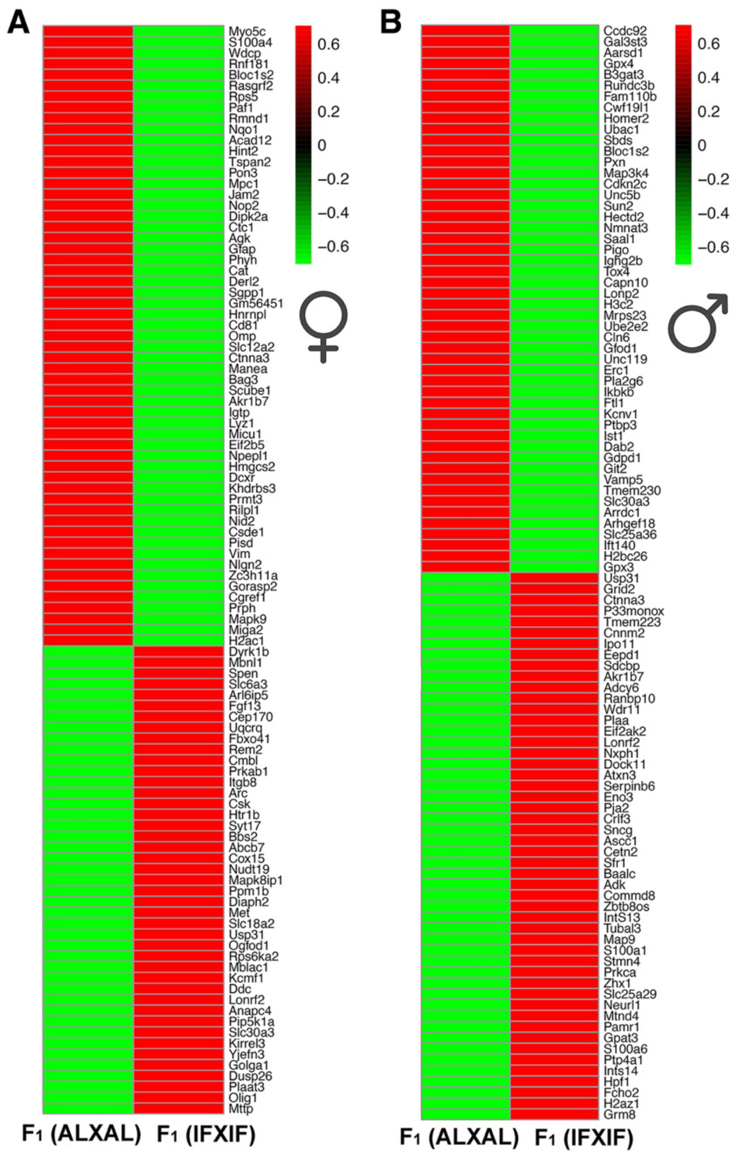


**Figure S13. Global proteomic heatmap analysis reveals extensive sensory-associated cortical protein remodeling in offspring derived from intermittent fasting parents.** (A) Heatmap showing representative top differentially expressed proteins in female offspring derived from intermittent fasting (IF) parents relative to offspring derived from ad libitum (AL) parents. Differentially expressed proteins included molecules associated with sensory signaling, neuronal communication, membrane transport, synaptic organization, and stress-response pathways, including *Slc12a2, Ctnna3, Rasgrf2, Myo5c, Nlgn2, Kirrel3, Prph, Fgf13,* and *Nqo1*. (B) Heatmap showing representative top differentially expressed proteins in male offspring derived from IF parents relative to controls. Differentially expressed proteins included molecules associated with neurotransmission, glutamatergic signaling, receptor trafficking, neuronal excitability, synaptic plasticity, and intracellular transport, including *Grm8, Erc1, Adcy6, Kcnv1, Pla2g6, Sncg, Map9, Dab2,* and *Wdr11*. These findings demonstrate persistent intergenerational remodeling of sensory-associated and neuronal signaling networks induced by parental IF exposure. Color scale represents relative protein expression levels.

**Supplementary Tables:**

Table S1: Differentially_Methylated_Genes_List_F0_Cortex_AL vs IF
Table S2: GO_enrich_hypo_F0_Cortex_AL vs IF
Table S3: GO_enrich_hyper_F0_Cortex_AL vs IF
Table S4: F0_Cortex_IFvsAL_DNA_RNA_common
Table S5: F0_Cortex_IFvsAL_DNA_Protein_common
Table S6: F0_17_common_gene
Table S7: F0_103_common_protein
Table S8: F1 Female DMR
Table S9: F1 Male DMR
Table S10: F1 Female GO_enrich_hypo
Table S11: F1 Female GO_enrich_hyper
Table S12: F1 Male GO_enrich_hypo
Table S13: F1 Male GO_enrich_hyper
Table S14: F1 Female DEG
Table S15: F1 Male DEG
Table S16: F1 Female DEP
Table S17: F1 Male DEP
Table S18: F1 (ALXAL) vs F1 (IFXIF)_Female_59_DMR_DEP Common Proteins
Table S19: F1 (ALXAL) vs F1 (IFXIF)_Male_59_DMR_DEP Common Proteins
Table S20: F0_F1 Male_F1 Female_DMR
Table S21: F0_F1 Male_F1 Female_DEG
Table S22: F0_F1 Female_DEP
Table S23: F0_F1 Male_DEP
